# Development of MAPT S305 mutation models exhibiting elevated 4R tau expression, resulting in altered neuronal and astrocytic function

**DOI:** 10.1101/2023.06.02.543224

**Authors:** KR Bowles, DA Pugh, C Pedicone, L Oja, SA Weitzman, Y Liu, JL Chen, MD Disney, AM Goate

## Abstract

Due to the importance of 4R tau in the pathogenicity of primary tauopathies, it has been challenging to model these diseases in iPSC-derived neurons, which express very low levels of 4R tau. To address this problem we have developed a panel of isogenic iPSC lines carrying the *MAPT* splice-site mutations S305S, S305I or S305N, derived from four different donors. All three mutations significantly increased the proportion of 4R tau expression in iPSC-neurons and astrocytes, with up to 80% 4R transcripts in S305N neurons from as early as 4 weeks of differentiation. Transcriptomic and functional analyses of S305 mutant neurons revealed shared disruption in glutamate signaling and synaptic maturity, but divergent effects on mitochondrial bioenergetics. In iPSC-astrocytes, S305 mutations induced lysosomal disruption and inflammation and exacerbated internalization of exogenous tau that may be a precursor to the glial pathologies observed in many tauopathies. In conclusion, we present a novel panel of human iPSC lines that express unprecedented levels of 4R tau in neurons and astrocytes. These lines recapitulate previously characterized tauopathy-relevant phenotypes, but also highlight functional differences between the wild type 4R and mutant 4R proteins. We also highlight the functional importance of *MAPT* expression in astrocytes. These lines will be highly beneficial to tauopathy researchers enabling a more complete understanding of the pathogenic mechanisms underlying 4R tauopathies across different cell types.

## INTRODUCTION

The *MAPT* gene, encoding the protein tau, is alternatively spliced at its N- and C-terminal ends, resulting in the expression of 6 different isoforms in the human brain. These can be categorized into two groups, dependent on the inclusion or exclusion of Exon 10; inclusion results in the expression of 4 microtubule binding repeat domains (4R tau), whereas exclusion results in only 3 (3R tau). The 4R:3R tau ratio in healthy adult human brain is roughly 1:1^1–3^, while expression in the developing fetal brain is almost exclusively 3R tau^4^. Several primary tauopathies, including progressive supranuclear palsy (PSP), corticobasal degeneration (CBD) and some forms of frontotemporal lobar dementia-*MAPT* (FTLD-*MAPT*) are considered “4R tauopathies” due to the specific accumulation of this tau isoform in neuronal and astrocytic inclusions^5^. The functional impact of aberrantly increased 4R tau expression and accumulation on cellular function is not fully understood, although these processes are likely to be key for determining mechanisms of disease pathogenesis and subsequent therapeutic intervention.

Human iPSC models have proven themselves a valuable tool for the study of early cellular perturbations and mechanisms contributing to neurodegenerative disorders: for example, both 2D neuronal monocultures and 3D organoid models have recapitulated key phenotypes associated with tauopathies, and reveal unique phenotypes of interest^6–17^. However, these models, like other iPSC-derived neuronal models, most closely resemble fetal brain^18^, and therefore express almost exclusively 3R tau. Modeling the functional effect of aberrantly expressed 4R tau and pathogenic mechanisms that underlie 4R tauopathies has therefore been challenging in these systems.

To date, there have been numerous efforts to increase the expression of 4R tau in iPSC models, with mixed results^19–23^. However, these models require extended periods in culture or complex culturing techniques and manipulations to achieve reasonable expression of 4R tau^19–21^. While neurons derived from directly differentiated human fibroblasts more closely recapitulate the epigenetic signature of aging and express 4R tau at a level equivalent to adult human brain^22, 23^, this model is limited in its capacity for expansion, isogenic correction and accessibility to the wider scientific field.

To address this issue, we have developed an isogenic set of iPSC lines with autosomal dominant S305 mutations, which cause FTLD-*MAPT*^24–27^. S305 is the last codon of *MAPT* Exon 10, and is therefore part of the RNA hairpin loop structure that regulates exon 10 alternative splicing^28^. Three mutations have been identified in this codon: two missense mutations (S305N, S305I) and one synonymous mutation (S305S) that all alter *MAPT* splicing, increasing Exon 10 inclusion. However, while each clinical case has been described as a variant of frontotemporal dementia (FTD), each mutation is varied in its clinical and neuropathological presentation^24–27^.

Here, we present a novel isogenic panel of iPSCs carrying S305N, S305I or S305S mutations, derived from four independent donors. Each isogenic series includes WT, heterozygous and homozygous mutations. We demonstrate that these mutations significantly increase 4R tau expression in iPSC-neurons and astrocytes from as early as 4 weeks of differentiation. Furthermore, we have characterized both shared and divergent cellular phenotypes associated with increased 4R tau between missense and synonymous splicing mutations, converging on synaptic function and mitochondrial metabolism in neurons, and phagocytic function and inflammatory state in astrocytes. These lines are a valuable and necessary tool for progressing tauopathy research, and are freely available to the scientific community.

## RESULTS

### Generation of a novel isogenic panel of *MAPT* S305 mutation iPSC lines

iPSC-derived models of tauopathy are notoriously imperfect due to low expression of *MAPT* exon 10 and consequently result in the absence of detectable 4R tau. To address this, we generated iPSC lines with *MAPT* S305 splice-site mutations (Figure 1A-B), in order to increase Exon 10 inclusion in these models. We identified peripheral blood monocyte cells (PBMCs) from individuals heterozygous for *MAPT* S305I and S305N mutations as part of the ALLFTD study, as well as lymphoblastoid cell lines (LCLs) from an S305S mutation carrier at the Sydney Brain Bank (SBB, Sydney, Australia) (Table 1). PBMCs and LCLs were reprogrammed to iPSC using Sendai virus mediated gene transfer of *OCT4*, *SOX2*, *KFL4* and c-*MYC* at the Icahn School of Medicine at Mount Sinai Stem Cell core facility. Following reprogramming, the mutation status, karyotypic normality and expression of pluripotency markers of one clone per donor were confirmed and selected for CRISPR/Cas9 genome editing (Figure 1C-D).

**Figure 1.**
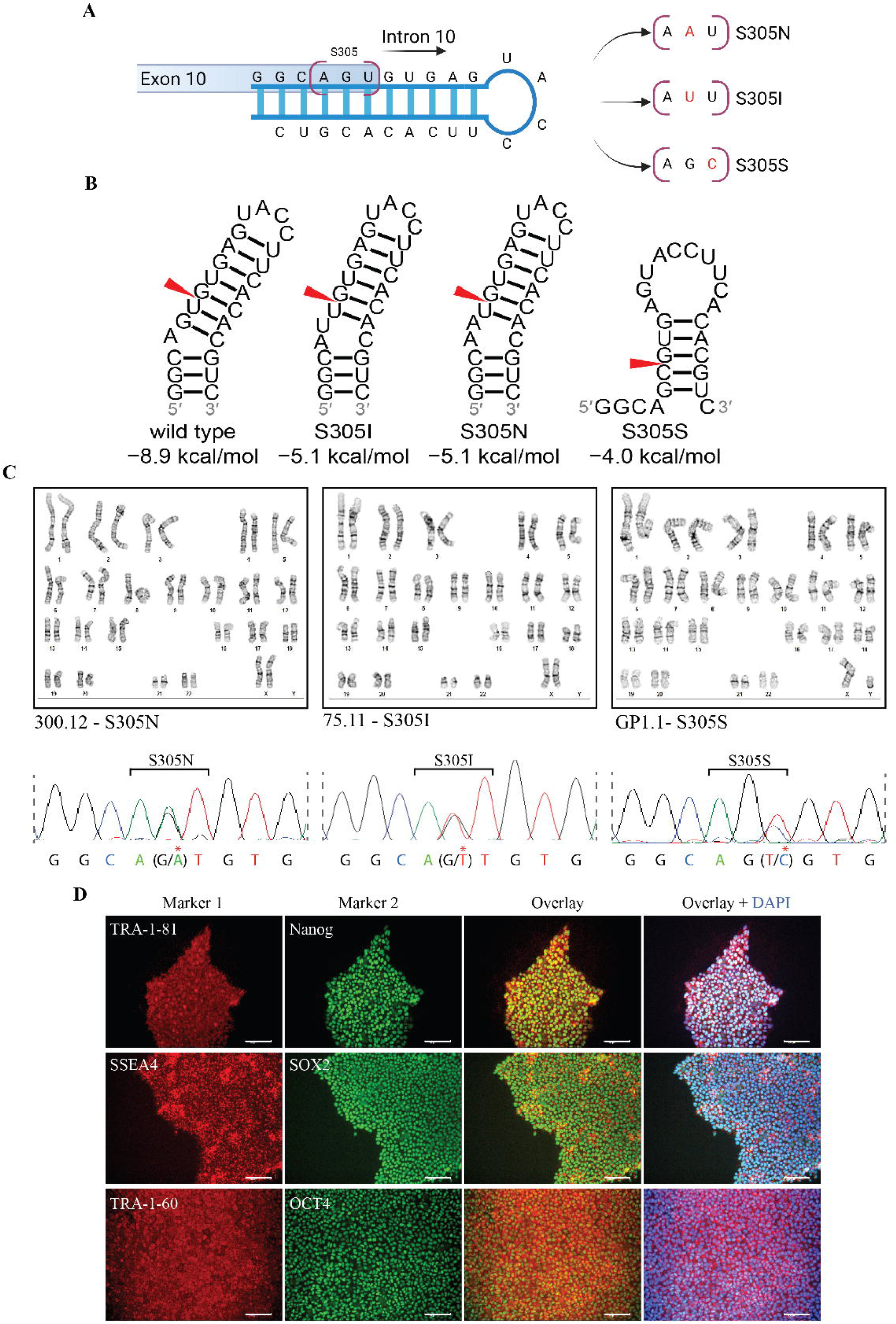
Generation of *MAPT* S305 mutation iPSC lines. **A.** Schematic of the *MAPT* exon 10 – intron 10 RNA hairpin, highlighting the location of S305 and the sequence modifications that define each mutation identified at this codon. **B.** Predicted structures and free energies (kcal) of the Exon 10 – Intron 10 splice site hairpin for WT and S305 mutation sequences. Modifications are indicated with a red arrow. Structure predictions were carried out using RNAstructure^91^. **C.** G-band karyotyping of reprogrammed parent clones selected for CRISPR editing from each *MAPT* mutation donor, paired with sequencing traces confirming heterozygote S305 mutation status. Red asterisks denote the mutant variant. **D.** Representative example of immunofluorescence of pluripotency markers in reprogrammed iPSC clones. Scale bar = 100µm.

**Table 1.** Description of isogenic sets of *MAPT* S305 mutation iPSC lines

| Parent ID | Clone ID | Donor Genotype | Isogenic Genotype | Description | Initial cell type | Source repository | iPSC available from | Sex | PAM edit |
| --- | --- | --- | --- | --- | --- | --- | --- | --- | --- |
| 75.11 | - | WT/S305I | - | Reprogrammed clone 3 | PBMC | NCRAD | NCRAD | XX | - |
| 75.11 | IW1A12 | WT/S305I | WT/WT | Corrected control |  |  | NCRAD | XX | Het |
| 75.11 | IW1B5 | WT/S305I | WT/WT | Corrected control |  |  | NCRAD | XX | None |
| 75.11 | IW1F3 | WT/S305I | WT/WT | Corrected control |  |  | NCRAD | XX | None |
| 75.11 | IW1G8 | WT/S305I | WT/WT | Corrected control |  |  | NCRAD | XX | None |
| 75.11 | IH1A8 | WT/S305I | WT/S305I | Unedited clone |  |  | NCRAD | XX | None |
| 75.11 | IH1A5 | WT/S305I | WT/S305I | Unedited clone |  |  | NCRAD | XX | None |
| 75.11 | IH1C10 | WT/S305I | WT/S305I | Unedited clone |  |  | NCRAD | XX | None |
| 75.11 | IH1B9 | WT/S305I | S305I/S305I | Homozygous edit |  |  | NCRAD | XX | Hom |
| 75.11 | IH1A4 | WT/S305I | S305I/S305I | Homozygous edit |  |  | NCRAD | XX | Hom |
| 300.12 | - | WT/S305N | - | Reprogrammed clone 3 | PBMC | NCRAD | NCRAD | XX | - |
| 300.12 | NWBB7 | WT/S305N | WT/WT | Corrected control |  |  | NCRAD | XX | Het |
| 300.12 | NW1B10 | WT/S305N | WT/S305N | Unedited clone |  |  | NCRAD | XX | None |
| 300.12 | NW1C7 | WT/S305N | WT/S305N | Unedited clone |  |  | NCRAD | XX | None |
| 300.12 | NH1E7 | WT/S305N | WT/S305N | Unedited clone |  |  | NCRAD | XX | Het |
| 300.12 | HBE10 | WT/S305N | S305N/S305N | Homozygous edit |  |  | NCRAD | XX | None |
| 300.12 | HCB8 | WT/S305N | S305N/S305N | Homozygous edit |  |  | NCRAD | XX | Hom |
| 300.12 | HAE11 | WT/S305N | S305N/S305N | Homozygous edit |  |  | NCRAD | XX | None |
| GP1.1 | - | WT/S305S | - | Reprogrammed clone 3 | Lymphoblast | SBB | NeuraCell | XY | - |
| GP1.1 | WAC11 | WT/S305S | WT/WT | Corrected control |  |  | NeuraCell | XY | Hom |
| GP1.1 | SW1B4 | WT/S305S | WT/S305S | Unedited clone |  |  | NeuraCell | XY | None |
| GP1.1 | SW1D8 | WT/S305S | WT/S305S | Unedited clone |  |  | NeuraCell | XY | None |
| GP1.1 | SH1F6 | WT/S305S | S305S/S305S | Homozygous edit |  |  | NeuraCell | XY | Hom |
| GP1.1 | SH1G8 | WT/S305S | S305S/S305S | Homozygous edit |  |  | NeuraCell | XY | Hom |
| F13505.1 | - | WT/WT | - | Parent line | Fibroblast | WashU ADRC | NeuraCell | XX | - |
| F13505.1 | S2A3 | WT/WT | WT/WT | Unedited clone |  |  | NeuraCell | XX | None |
| F13505.1 | I1E12 | WT/WT | WT/WT | Unedited clone |  |  | NeuraCell | XX | None |
| F13505.1 | NEA11 | WT/WT | WT/WT | Unedited clone |  |  | NeuraCell | XX | Hom |
| F13505.1 | I1B10 | WT/WT | WT/S305I | Heterozygous edit |  |  | NeuraCell | XX | Het |
| F13505.1 | NDG11 | WT/WT | WT/S305N | Heterozygous edit |  |  | NeuraCell | XX | Het |
| F13505.1 | S3H5 | WT/WT | WT/S305S | Heterozygous edit |  |  | NeuraCell | XX | Het |

As donor and clonal variability are major sources of inconsistency in iPSC phenotypic characterization^29^, it is important to generate isogenic corrected controls from donor lines to isolate effects specific to the mutation of interest. To this end, each mutation donor line underwent CRISPR/Cas9-mediated correction to wild type (WT) (Table 1, Table S1, Figure S1). In addition, we hypothesized that homozygous mutations would likely exhibit increased *MAPT* Exon 10 inclusion and more severe phenotypes than heterozygous mutants. We therefore also edited each donor line to be homozygous for their original mutation (Table 1, Figure S1). In order to control for clonal variability and selective pressures from the editing process, additional unedited clones per donor line were retained (Table 1, Figure S1). An additional WT iPSC line derived from a cognitively normal individual was selected for CRISPR/Cas9 induction of each S305 mutation onto the same control background (Table 1, Table S1, Figure S1), in order to confirm that downstream functional effects were a result of *MAPT* mutation rather than genetic backgrounds of each donor. This process resulted in a unique panel of 31 iPSC lines, derived from four donors, which are heterozygous or homozygous for *MAPT* S305S, S305I or S305N splicing mutations (Table 1). Four of these clones were previously reported as a resource available from the Tau Consortium^30^, and are now included in the wider panel of S305 mutation lines reported here. All lines are freely available upon request from the National Centralized Repository for Alzheimer’s disease (NCRAD; https://ncrad.iu.edu/) or the Tau Consortium via the Neural Stem Cell Institute (NSCI; http://neuralsci.org/tau).

### *MAPT* S305 mutation neurons express 4R tau as early as 4 weeks of differentiation

To date, iPSC-neuron models of tauopathy have been unable to fully model the effect of increased 4R *MAPT* expression or 4R tau accumulation as observed in PSP and FTD brain, due to their similarity to fetal-age cells that express primarily the 3R isoform^18–20^. *MAPT* S305 mutations are predicted to increase Exon 10 inclusion due to destabilization of the Exon 10 – Intron 10 RNA hairpin loop (Figure 1B), and as such are neuropathologically associated with the accumulation of 4R tau^24–27^. We therefore predicted that this hairpin loop destabilization would facilitate higher 4R tau expression in iPSC models.

We carried out forebrain neuron directed differentiation from our panel of S305 mutation iPSC lines and isogenic controls, and assayed Exon 10 inclusion at 4, 6 and 8 weeks of differentiation (Figure 2A-D, Figure S2A-D). Total *MAPT* expression and Exon 10 percent spliced in (PSI) values were assessed from RNA-seq data (Figure 2A, Figure S2A). Interestingly, there was a significant effect of weeks of differentiation (*p* = 0.016) on total *MAPT* expression when correcting for parent line variability, with expression tending to reduce in mutant neurons with time (Figure S2A). This effect remained significantly significant in only S305N homozygote lines at 8 weeks of differentiation following post-hoc testing (*p* = 0.022). In contrast, there was a significant effect of weeks of differentiation on Exon 10 PSI in WT neurons (F^(5, 18)^= 3.1, *p* = 0.03), with increased inclusion over time, as has been previously observed^19^ (Figure 2A). Surprisingly, there was no effect of weeks of differentiation on the inclusion of Exon 10 in S305S, S305I or S305N neurons, indicating that these mutations significantly increase 4R inclusion from as early as 4 weeks of differentiation, and remain stable over time (Figure 2A, Figure S2B). S305N mutations resulted in the largest increase in Exon 10 PSI, with a clear gene dosage effect contributing to the proportion of Exon 10 inclusion (mean PSI WT/S305N ∼ 0.3-0.4, S305N/S305N ∼ 0.6, Figure 2A). S305I mutations resulted in the second highest increase in Exon 10 PSI, although due to high variability in heterozygous clones, this was only statistically significant in homozygous mutants, despite no clear gene dosage effect (both WT/S305I and S305I/S305I mean PSI ∼ 0.1-0.16). Increased Exon 10 inclusion was confirmed by qRTPCR in S305I lines compared to isogenic controls across time (Figure S2B). S305S mutations had the mildest effect on Exon 10 PSI (Figure 2A, Figure S2B), and there was no mutation dosage effect (WT/S305S and S305S/S305S mean PSIs ∼ 0.06-0.08, Figure 2A). This was surprising to us, as predictions of RNA stability estimated S305S to most greatly destabilize hairpin structure, and S305N and S305I were estimated to have similar disruption. Regardless, all mutations influenced Exon 10 splicing.

**Figure 2.**
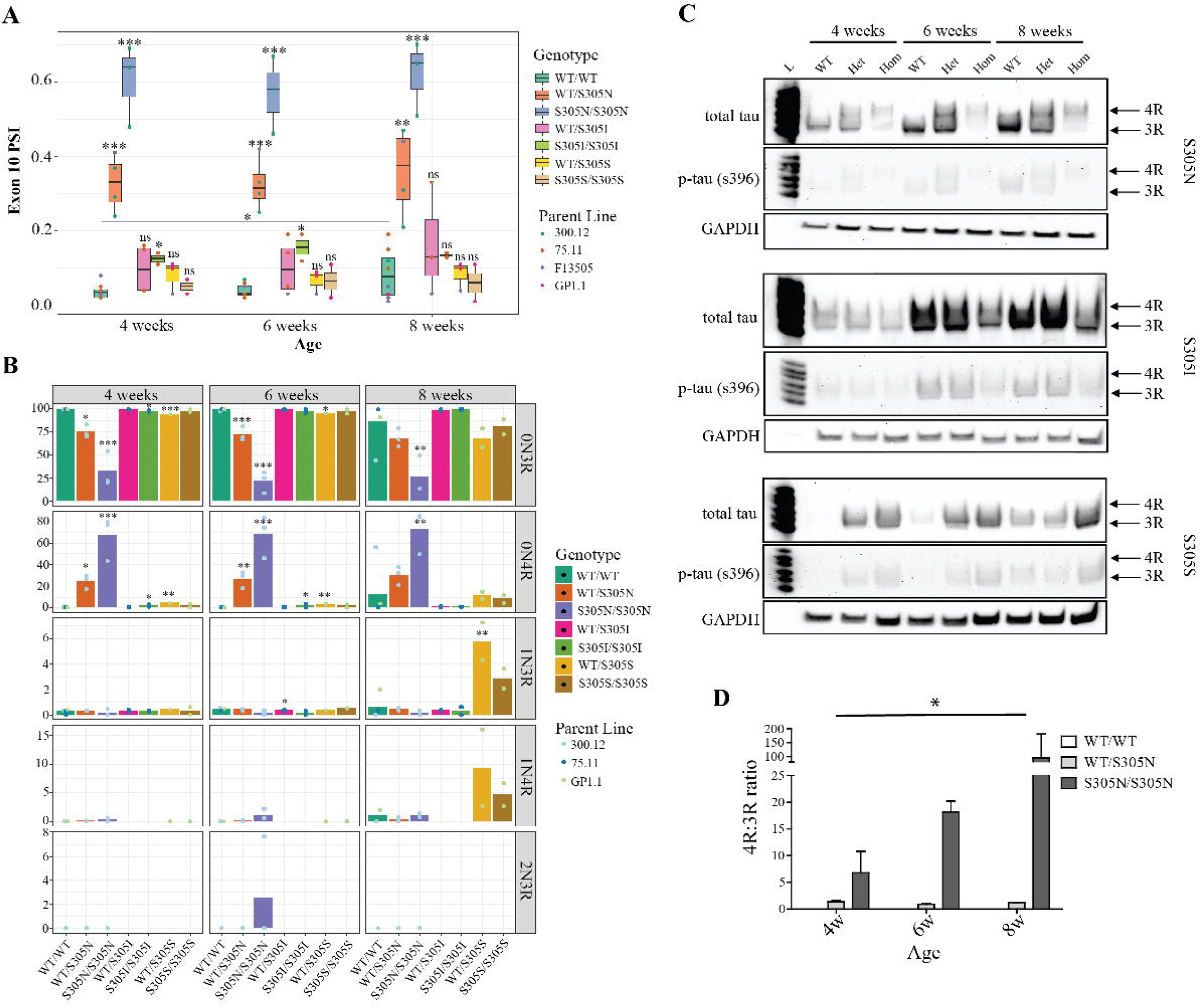
*MAPT* S305 mutations significantly increase Exon 10 inclusion in iPSC-neurons. **A**. Percent spliced in (PSI) values for *MAPT* Exon 10 derived from RNA-seq data of 4, 6 and 8 week old iPSC-neurons per clone, separated by mutation. Error bars = SEM. N = 2-8. One-way ANOVA at each time point including parent line as a covariate, Tukey post-hoc testing compared to isogenic controls. **B.** Targeted *MAPT* ISOseq for 5 main neuronally expressed transcripts for each clone. Error bars = SEM. N = 2-8. Two-way ANOVA, Tukey post-hoc testing compared to isogenic controls. **C.** Representative western blot for total tau and phosphorylated tau (p-tau (S396)) in neurons derived from one clone per genotype over time. L = tau ladder. GAPDH used as a loading control. **D.** Quantification of 4R:3R ratio for S305N clones, represented in D. Error bars = SEM. Two-way repeated measures ANOVA with Bonferroni post-hoc testing. *p < 0.05, **p < 0.01, ***p < 0.001

In order to investigate whether S305 splicing mutations may also destabilize regulation of *MAPT* N-terminal splicing, and to characterize the expression of full-length isoforms, we carried out targeted ISOseq on the same S305 mutation and isogenic control iPSC-neurons (Figure 2B). As expected, iPSC-neurons expressed primarily 0N isoforms, however S305N mutants significantly reduced the proportion of 0N3R at all three time points (WT mean = 86-99%, WT/S305N mean = 68-75%, S305N/S305N mean = 21.5-32%) in favour of significantly increased 0N4R (WT = 0.3-12%, WT/S305N = 24-30%, S305N/S305N = 67-73%) (Figure 2B). Similar to the short-read sequencing data, the effect of S305I and S305S mutations on Exon 10 inclusion were much milder, although both mutations expressed significantly higher levels of 0N4R at 4- and 6-weeks of differentiation compared to isogenic controls (S305I = 0.7-2%, S305S = 3.4-11.5%, Figure 2B). Interestingly, there was a slight increase in the detection of 1N isoforms in S305S and S305N mutations, with a significant increase in 1N3R in S305S heterozygotes at 8-weeks (WT = 0.6%, WT/S305S = 5.8%, S305S/S305S = 2.8%, Figure 2B). S305N mutations showed a trend towards increased 1N4R over time, although this did not reach statistical significance, likely due to very low levels of detection (Figure 2B).

To validate the transcriptomic data and confirm that we could observe 4R tau protein in S305 mutation neurons, we carried out western blots (Figure 2C-D, Figure S2C-D). There was a significant effect of both time and genotype on 4R tau expression in S305N neurons (F_(2,18)_ = 3.63, *p* < 0.05). We observed a clear increase in 4R expression in S305N neurons compared to isogenic controls, with homozygous lines expressing almost exclusively 4R tau, as compared to heterozygous mutants that had roughly 1:1 3R:4R tau (Figure 2C-D). In comparison, we were unable to detect 4R tau by western blot in S305I and S305S mutation lines, likely due to relatively low expression of 4R compared to S305N mutants (Figure 2C). In contrast to previous reports for other *MAPT* mutations^6, 8^, we did not observe accumulation of tau nor mutant tau over time for any S305 mutation (Figure 2C, Figure S2C). Instead, there was a trend towards reduced total tau protein in S305N and S305S mutants, although this did not reach significance (Figure 2C, Figure S2C). Despite potentially lower tau levels in S305N and S305S mutant neurons, we did observe a significantly increased phosphorylated tau on serine 396 (p-tau_s396_) to total tau ratio (p-tau/total tau) in homozygous mutants at each time point compared to isogenic controls (Figure 2C, Figure S2D).

### Different *MAPT* S305 mutations have both distinct and shared transcriptional effects

In contrast to the synonymous S305S splicing mutation, non-synonymous S305N and S305I mutations both increase 4R *MAPT* expression and alter the tau protein sequence. We hypothesized that expression of mutant 4R tau in these lines may therefore have differential effects on neuronal function compared to expression of WT 4R tau. We therefore carried out Gene Set Enrichment Analysis (GSEA) and Ingenuity pathway analysis (IPA) on our RNA-seq data to identify and characterize the shared and divergent functional effects of each S305 mutation in iPSC-neurons.

When comparing the number of significantly enriched pathways by GSEA shared across all three S305 mutations, we find that heterozygous mutations shared 151 (7.3%) significantly enriched pathways at 4 weeks, but lines became more divergent with age, sharing only 3 pathways (0.3%) by 8 weeks (Figure S2E). In contrast, homozygous mutations retained a consistently similar proportion of overlap across time (23-26 pathways, 1.4-1.9%) (Figure S2F). Inspection of the top 20 significant pathways for each gene dosage at each time point revealed the pathways most strongly impacted by each mutation (Figure 3, Table S2). The S305N mutation was associated with strong downregulation of neurotransmitter release and glutamate signaling, the severity of which correlated with mutant allele dosage (Figure 3, Table S2). In contrast, there was early upregulation of translation and ribosomal pathways at 4 weeks of differentiation, which lost significance with age (Figure 3, Table S2). Other pathways upregulated across differentiation were involved in cell cycle regulation and endoplasmic reticulum stress (Figure 3, Table S2).

**Figure 3.**
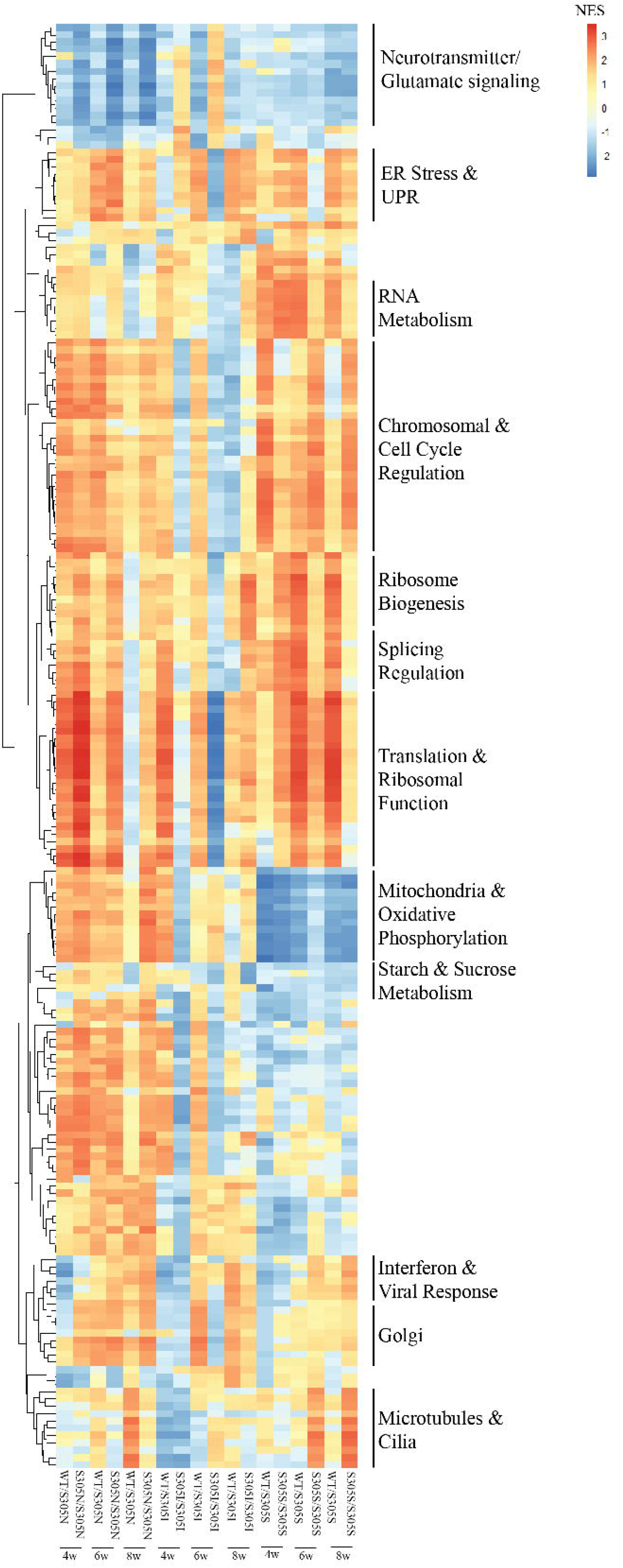
Shared and divergent transcriptomic effects of *MAPT* S305 mutations. Integration of the top 20 significantly enriched GSEA pathways across each genotype and age. Colour represents the normalized enrichment score (NES); red = predicted upregulation, down = predicted downregulation. Pathways underwent unsupervised clustering based on NES across samples and clusters manually annotated by function.

S305I mutation lines showed a more complex relationship between heterozygous and homozygous mutation carriers over the course of differentiation, but exhibited a similar pathway enrichment by 8 weeks (Figure 3, Table S2). This mutation was associated with progressive upregulation of interferon-related pathways, as well as increased viral response pathways (Figure 3, Table S2). Similar to S305N lines, we also observed upregulation of ribosomal and ER stress pathways, as well as responses to unfolded protein and incorrect protein responses (Figure 3, Table S2).

Finally, S305S mutation lines were also associated with upregulation of ribosomal and translational regulation pathways, increased ER and UPR pathways, as well as a time-dependent increase in interferon-related pathways, similar to S305N and S305I mutations (Figure 3, Table S2). Specific to S305S was downregulation of pathways associated with starch and sucrose metabolism, including glycolysis and alcohol and sterol biosynthesis (Figure 3, Table S2).

### *MAPT* S305 mutations accelerate synaptic maturation and neuronal hyperexcitability

When comparing the top GSEA pathways across all S305 mutations, we observed downregulation of pathways associated with neurotransmitter signaling and glutamate receptor signaling, although the effect of this suppression was greater in S305N lines (Figures 3, 4A). Downregulation of these pathways was surprising to us, as we previously characterized upregulation of glutamatergic receptor genes in *MAPT* V337M mutation neurons^6^, and other *MAPT* mutation iPSC-neurons have been found to exhibit electrophysiological hyperexcitability^7^. However, further inspection of genes involved in these pathways using Ingenuity Pathway Analysis (IPA) indicated *GABAergic signaling* as the most significantly enriched canonical pathway amongst this gene set (overlap = 95/132 genes (72%), *p* = 6.21x10^-118^), suggesting a potential loss of inhibitory regulation in S305 neurons.

We then focused on heterozygote mutant carriers only, because two parent lines contributed to these signals compared to only one parental donor for the homozygote edits. Here, we observed a differential mutation effect; S305S neurons showed upregulation of LTP and synaptic signaling pathways early in differentiation, whereas these pathways were suppressed in favour of upregulated seizure-related pathways in S305N neurons, with S305I neurons landing in between the two (Figure 4B). This pattern was consistent with the proportion of Exon 10 expressed by each mutation, and may reflect a transition in neuronal synaptic dysregulation with increasing Exon 10/4R tau accumulation over time, i.e. early increased excitatory signaling followed by downregulation of inhibitory signaling.

**Figure 4.**
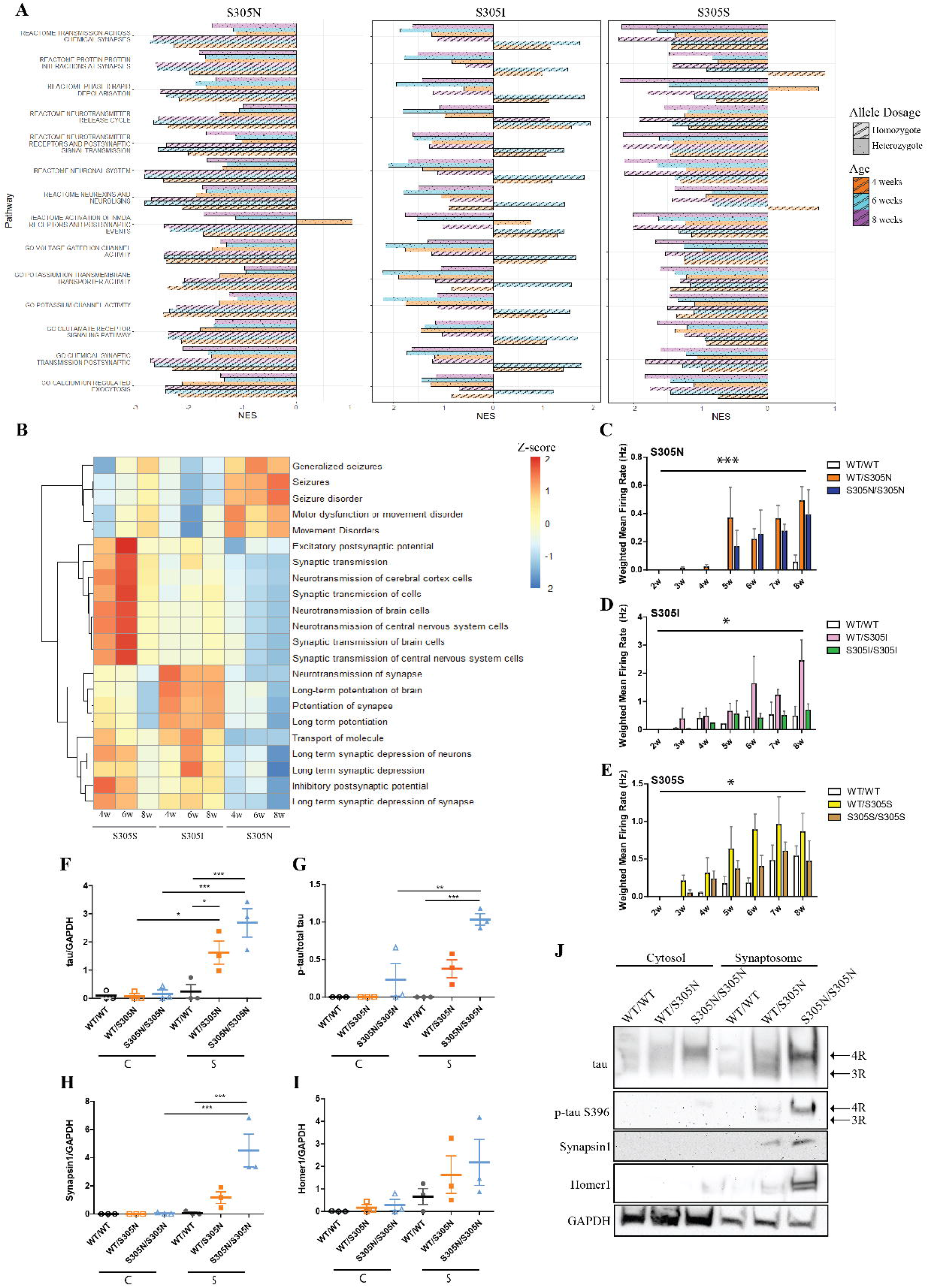
Neurons with *MAPT* S305 mutations have accelerated synaptic maturity and are electrophysiologically active earlier than isogenic controls. **A.** Normalized enrichment score (NES) of synaptic-related pathways in ***Figure 3*** for each S305 mutation across time. **B.** Z-scores for enriched pathways from IPA analysis of genes derived from ***A*** for each heterozygous mutation at each time point. **C-E.** Weighted mean firing rate (Hz) derived from multi-electrode array analysis of isogenic sets of ***C.*** S305N, ***D.*** S305I and ***E.*** S305S iPSC-neurons every week from 3-8 weeks of age. Error bars = SEM. N = 3 independent differentiations, averaged from 3 technical replicates per genotype/clone. Two-way ANOVA with Bonferroni post-hoc testing. **F-I.** Quantification of western blots for ***F.*** total tau, ***G.*** p-tau (S396), ***H.*** Synapsin1 and ***I.*** Homer1, normalized to GAPDH loading control in 6-week old S305N mutation cytosol (C) and synaptosome (S) fractions compared to isogenic controls, as represented in ***J.*** Error bars = SEM. N = 3. One way ANOVA with Bonferroni post-hoc testing. *p < 0.05, **p < 0.01, ***p < 0.001 **J.** Representative western blot of cytosolic and synaptosome fractions derived from 6-week old S305N mutation neurons and isogenic controls, labelled for total tau and p-tau (S396), as well as synaptic markers Synapsin1 and Homer1. GAPDH used as loading control.

To determine whether altered gene expression pathways translated to functional changes in neuronal firing we carried out multielectrode array (MEA) analysis of S305 mutation iPSC-neurons. All three mutations exhibited an early and increased number of spikes and significantly increased weighted mean firing rate compared to isogenic controls, from as early as 3 weeks of differentiation (Figure 4C-E, Figure S3A), consistent with previous reports of early hyperexcitability in *MAPT* mutation neurons^7^. Interestingly, there was no mutation dosage effect in S305N neurons, despite differences in 4R protein expression, although both WT/S305N and S305N/S305N edits were more active than the isogenic controls from 3 weeks of differentiation (Figure 4C, Figure S3A). S305I neurons also showed increased firing from as early as 3 weeks, although curiously the heterozygote mutants were more active than the homozygotes (Figure 4D, S3A), which was consistent with the pattern of pathway enrichment from the GSEA analysis (Figure 4A). Finally, S305S mutants exhibited the same hyperexcitability from 3 weeks, despite much lower levels of 4R tau compared to S305N mutants (Figure 4E, S3A). The isogenic control line showed a similar level of spontaneous activity by 7-8 weeks (Figure 4E, S3A), consistent with the pattern of Exon 10 inclusion we observed across this time-frame (Figure 2A, B).

Given the transcriptomic and functional effects on synaptic firing observed across S305 mutations, and that mutant tau protein has been reported to accumulate in and interact with components of the synapse^31, 32^, we isolated synaptosomes from S305 mutation iPSC-neurons and isogenic controls to assess tau accumulation and synaptic maturity (Figure 4F-J, Figure S3B-F). Consistent with previous reports from mouse and human brain^33–35^, we found that total tau and p-tau_s396_ preferentially accumulated in S305N and S305I synaptosomes compared to the cytosol, and in comparison to isogenic controls (Figure 4F-G,J, Figure S3B-C), although S305S lines had similar total tau and p-tau_s396_ levels to controls (Figure S3D-E). We also saw increased presence of synaptic markers Homer1 and Synapsin1 in S305I and S305N synaptosome preparations compared to isogenic controls, and a trend towards increased synaptic marker expression in S305S synaptosomes (Figure 4H-J, S3B-E). This suggests that synapses may be more readily formed and/or matured in the presence of 4R tau, which may underlie the increased neuronal firing observed by MEA.

### Mitochondrial function is differentially affected by WT and mutant 4R tau

One of the most striking differences between S305 mutation lines by GSEA analysis was the effect on mitochondria-related pathways, such as *Oxidative Phosphorylation*, *Electron Transport Chain* and *Cellular Respiration* (Figure 3, Table S2). Curiously, these pathways were upregulated in both S305I and S305N neurons, but downregulated in S305S neurons, indicating a differential effect between increased WT 4R tau, and mutant 4R tau. IPA analysis confirmed the disparate effect of mutation on oxidative phosphorylation (Figure 5A).

**Figure 5.**
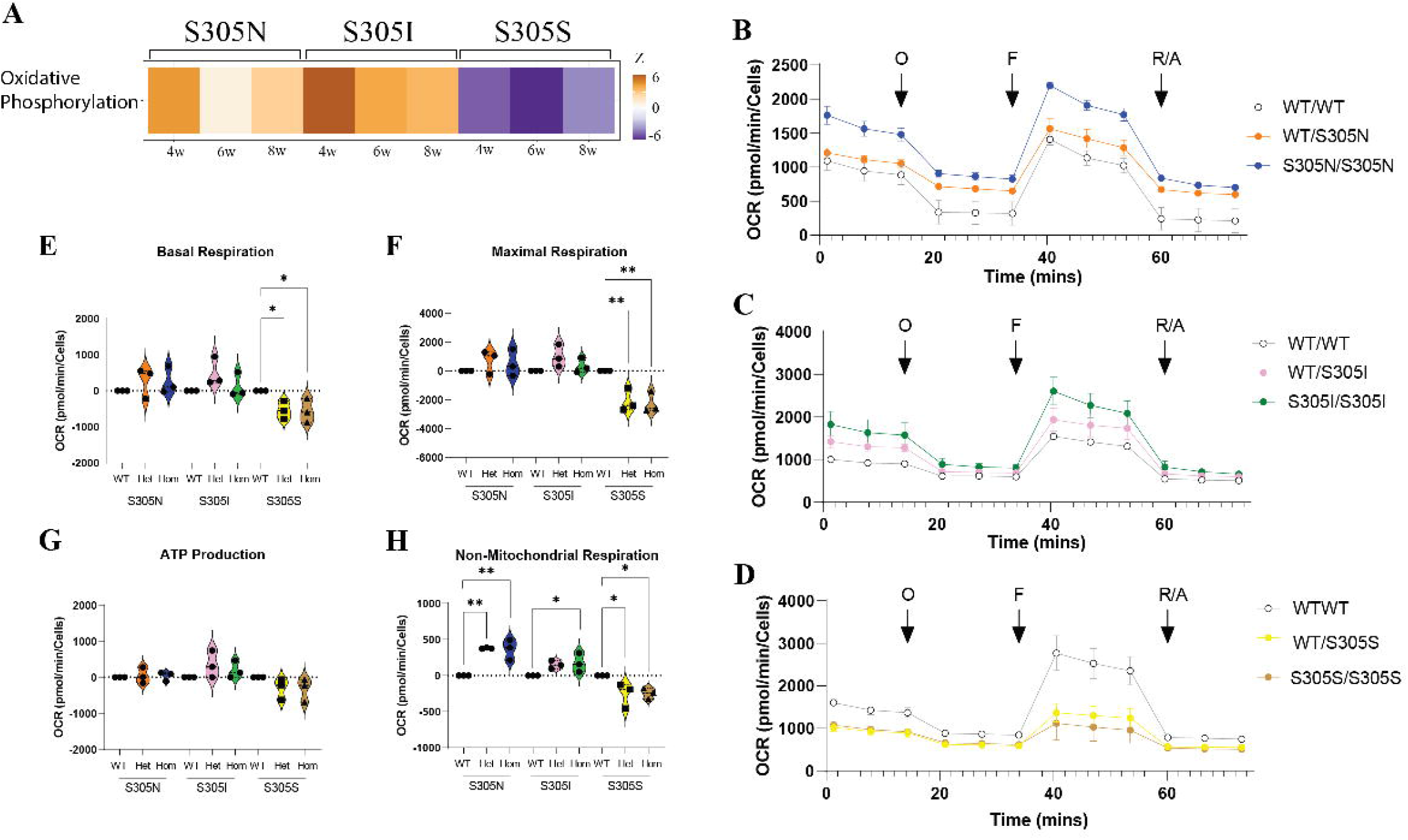
Mitochondrial bioenergetics are altered in *MAPT* S305 mutation neurons. **A.** Z-scores for enrichment of the Oxidative Phosphorylation pathway derived from IPA analysis of differentially expressed genes in heterozygous *MAPT* S305 mutation neurons compared to isogenic controls. Orange = predicted activation, purple = predicted suppression. **B-D.** Representative plots of oxygen consumption rate (OCR, pmol/min corrected for number of cells) in isogenic sets of ***B***. S305N, ***C.*** S305I and ***D.*** S305S neurons at 6 weeks, output from the Seahorse Mitochondrial stress test assay. Arrows indicate the time of treatment with mitochondrial stressors; O = Oligomycin, F = FCCP, RA = Rotenone/Antimycin-A. Error bars = SEM. **E-H.** Quantification of mitochondrial ***E.*** basal respiration, ***F.*** maximal respiration, ***G.*** ATP production and ***H.*** non-mitochondrial respiration relative to isogenic controls in *MAPT* S305 mutation neurons at 6 weeks. Error bars = SEM. N = 3 independent differentiations, averaged from 3 technical replicates each time. One-way ANOVA with Fisher’s LSD posthoc testing. *p < 0.05, **p < 0.01, ***p < 0.001

To functionally examine whether S305 mutations disrupt mitochondrial function, we then carried out Seahorse mitochondrial stress tests on 6-week old neurons (Figure 5B-H, S4A-B). Consistent with the pathway analyses, we observed a trend towards increased basal respiration in both S305N and S305I neurons compared to isogenic controls (Figure 5E), indicating more oxygen consumption was required to meet neuronal energy demands, but significant downregulation in S305S neurons (Figure 5E), suggesting reduced energy demand in these neurons. The same increased trend was present in the S305N and S305I neurons for both maximal respiration and spare respiratory capacity (Figure 5F, S4A), which may be reflective of their increased potential for hyperexcitability and therefore increased bioenergetic demand. Consistent with the RNA-seq data, these measurements were significantly reduced in S305S neurons (Figure 5F, S4A), potentially indicating reduced mitochondrial capacity for oxidative phosphorylation. There were no significant differences in either ATP production or proton leak between genotypes indicating that mitochondrial membranes were intact and sufficient ATP was being generated to meet energy demands in mutation neurons (Figure 5G, S4B). Interestingly, non-mitochondrial respiration was significantly increased in both S305N and S305I neurons (Figure 5H). High non-mitochondrial respiration has been associated with increased energy production via glycolysis, but has also been linked with increased inflammation and oxidative stress^36, 37^, both processes that have been associated with tauopathy^38–40^. In contrast, non-mitochondrial respiration was significantly reduced in S305S neurons (Figure 5H), indicating the majority of oxygen consumption in these cells was being used for mitochondrial respiration, and that cellular bioenergetics may be impaired.

### *MAPT* splicing is altered in S305 mutation astrocytes and is distinct from neuronal splicing regulation

Astrocytes express low levels of *MAPT*, which has been demonstrated to have significant effects on intrinsic astrocytic functions, including altered glutamate processing, neuronal support and amyloid-β-induced synaptotoxicity^41–43^. We therefore differentiated our S305 mutation panel into astrocytes, and assessed the effect of these mutations on astrocytic *MAPT* expression and splicing.

*MAPT* expression in astrocytes was around 50 fold lower than in neurons (neurons ∼10,000 reads per sample, astrocytes ∼200 reads per sample). Similar to the neurons, there was no significant effect of mutation on total *MAPT* expression after correction for donor, however there was a significant effect on Exon 10 PSI (F_9,15_ = 3.395, *p* < 0.05) (Figure 6A). Interestingly, S305I and S305S mutations had a larger impact on Exon 10 PSI in astrocytes compared to neurons, with a PSI of ∼20-30% in mutant astrocytes and 5-15% in mutant neurons (Figures 2, 6A). S305N mutations exhibited the most severe effect on splicing, with a clear mutation-dosage effect on Exon 10 PSI (Figure 6A).

**Figure 6.**
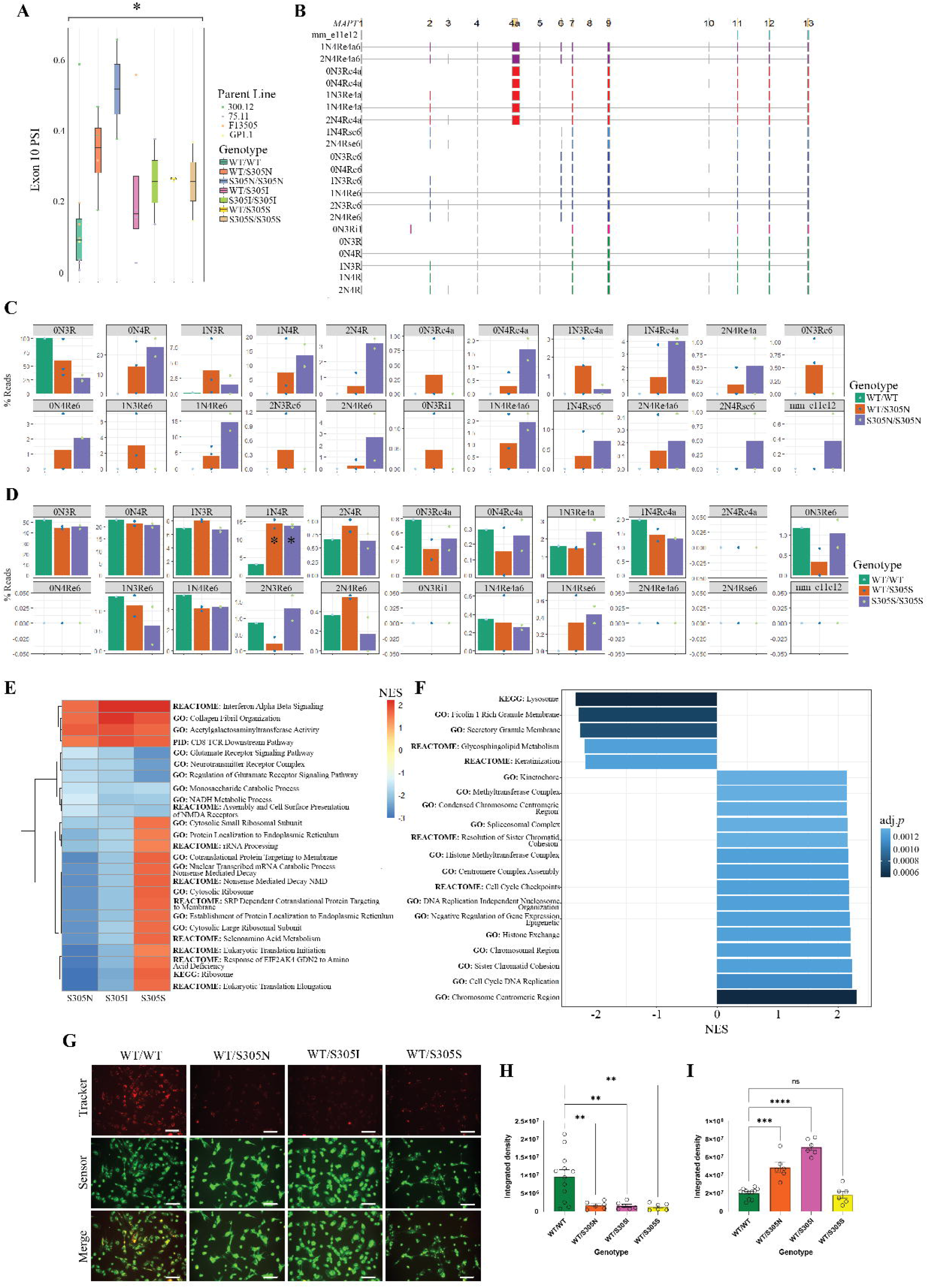
*MAPT* S305 astrocytes have a *MAPT* splicing profile distinct from neurons, and are associated with lysosomal dysfunction. **A.** Percent spliced in (PSI) values for *MAPT* Exon 10 derived from RNA-seq data in astrocytes per clone, separated by mutation. Error bars = SEM. N = 2-8. One-way ANOVA. **B.** Schematic of different *MAPT* transcripts observed in iPSC-astrocytes. “mm_e11e12” denotes most exons are missing, with the exception of 11 and 12. “e4a” indicates inclusion of Exon 4a, “6” or “e6” indicates inclusion of Exon 6. “se6” indicates inclusion of a smaller splice variant of Exon 6, and “i1” indicates retention of a portion of intron 1. Transcripts are coloured by their splice pattern (canonical = green, intron retention = pink, contains Exon 6 = dark blue, contains small Exon 6 = light blue, contains Exon 4a = red, contains Exons 4a and 6 = purple, mostly missing exons = pale blue). **C-D.** Percentage of reads mapping to each transcript identified from targeted ISOseq data in ***C.*** S305N and ***D.*** S305S mutation astrocytes. Isoform names are defined as described for ***B.*** Error bars = SEM. N = 1-3. One way ANOVA. E. Normalized enrichment scores (NES) for significantly enriched GSEA pathways shared across all three S305 mutation astrocyte lines. Blue = predicted downregulation, red = predicted upregulation. F. Normalized enrichment scores (NES) for significantly enriched GSEA pathways calculated as a function of Exon 10 PSI in S305 mutation astrocytes. Bar shading represents the adjusted enrichment *p*-value (adj.p) **G**. Representative images of Lysotracker (red) and Lysosensor (green) labelling in isogenic *MAPT* S305N, S305I and S305S mutation astrocytes compared to WT controls. Scale bar = 100µm. **H-I**. Quantification of integrated fluorescence density for ***H.*** Lysotracker and ***I.*** Lysosensor across all donor lines. Error bars = SEM. N = 6-12. One way ANOVA with Dunnett’s multiple comparisons tests. **p < 0.01, ***p < 0.001, ****p < 0.0001, ns = not significant.

Given the differences in Exon 10 inclusion compared to neurons, we carried out targeted *MAPT* ISOseq to examine differences in full-length transcripts between S305 mutations (Figure 6B-D). Technical issues with the second lane of sequencing resulted in very poor read depth and isoform detection for 75.11-derived and F13505-derived lines, therefore we focused these analyses on the most severe 300.12 (S305N) and milder GP1.1 (S305S) mutations. Interestingly, we identified a wider diversity of *MAPT* transcripts in astrocyte cultures compared to neurons (Figure 6B). We were able to detect both 1N and 2N transcripts in astrocytes, as well as the inclusion of Exon 6 and Exon 4a, which defines the “Big Tau” isoform that is present in the peripheral nervous system^44^ (Figure 6B). For the S305N isogenic corrected control line, only 0N3R and 1N3R were detected, whereas heterozygous and homozygous mutants expressed multiple other, primarily 4R transcripts (Figure 6C). Interestingly, S305S mutation astrocytes did not have significantly higher expression of 0N4R compared to their isogenic controls, but instead were enriched for 1N4R, and 1N4R including a portion of Exon 6 (1N4Rse6) (Figure 6D).

It should be noted that due to sequencing difficulties, these comparisons have only been conducted in one parent line per mutation, with a single isogenic corrected control. There was also significant variability between the two isogenic corrected controls, therefore it is not possible to confidently state the effect of S305 splicing mutations on the splicing of other exons in astrocytes. However, these data support the increased inclusion of Exon 10 in S305 mutation astrocytes, and demonstrate that astrocytes splice *MAPT* in a manner that is distinct from neurons.

### iPSC-astrocytes demonstrate cell-autonomous effects of mutant *MAPT* expression

In order to characterize the effect of S305 mutations and increased Exon 10 expression on astrocyte function, we carried out RNA-seq analysis on iPSC-astrocytes derived from our isogenic S305 mutation panel. We carried out GSEA analyses on differentially expressed genes within each mutation, while controlling for mutation dosage effects (Figure 6E, Table S3), and found 25 significantly enriched pathways shared across all three mutations. All three mutations resulted in upregulation of immune response-related pathways, as well as *collagen fibril organization* and *acetylgalactosaminyltransferase activity* (Figure 6E, Table S3), suggesting that mutant *MAPT* expression may be inducing an inflammatory phenotype in astrocytes, as well as altering glycosylation, which itself has been identified as a downstream effect of chronic inflammatory processes^36^. All three mutations were also associated with significant downregulation of glutamate and neurotransmitter signaling pathways (Figure 6E, Table S3), consistent with previous reports that *MAPT* mutation astrocytes are impaired in glutamate processing and clearance^41^.

In concordance with our observations in iPSC-neurons derived from the same lines, we observed disparate effects of synonymous vs. non-synonymous S305 mutations in the astrocytes (Figure 6E, Table S3). However, in contrast to their neuronal counterparts, S305N and S305I astrocytes were associated with significant downregulation of ribosomal and translation-regulation pathways, which were upregulated in neurons (Figure 3),while the same pathways were significantly upregulated in S305S astrocytes (Figure 6E). These data suggest that *MAPT* mutation and Exon 10 inclusion are likely to have disparate effects on similar cellular processes across different cell types.

We then repeated the GSEA analysis using the Exon 10 PSI value for each line as the defining variable, in order to assess the effect of Exon 10 inclusion on astrocyte gene expression (Figure 6F, Table S4). We found that increasing Exon 10 expression was associated with upregulation of multiple cell-cycle and chromosomal regulation pathways (Figure 6F), consistent with previous findings of abnormal cell-cycle re-entry in a humanized tau mouse model^45^. In contrast, increased *MAPT* Exon 10 PSI was significantly associated with downregulation of secretory membrane and lysosomal pathways (Figure 6F), indicating that in astrocytes, increased Exon 10 inclusion may be interfering with endocytic and exocytic processes.

We have previously observed lysosomal dysfunction in *MAPT* V337M mutation iPSC-organoids^6^, so we chose to validate this phenotype in S305 mutation astrocytes by labeling with lysosomal markers LysoTracker and LysoSensor. S305N, S305I and S305S mutant astrocytes all showed significantly reduced LysoTracker fluorescence intensity compared to isogenic controls, consistent with reduced lysosomal mass (Figure 6G-H). In contrast, LysoSensor fluorescence was significantly increased in S305 mutation astrocytes (Figure 6G-H), indicative of aberrant lysosomal acidification that would likely alter hydrolase activity and substrate degradation, which could be a precursor to pathogenic astrocytic tau accumulation in primary tauopathies.

### Tau internalization and inflammation are modulated by *MAPT* S305 mutations in astrocytes

Accumulation of tau in astrocytic tufts is a neuropathological feature of PSP and is commonly observed in FTD brain^46^. However, as *MAPT* expression is very low in these cells compared to neurons, the source of accumulated tau is thought to be exogenous to astrocytes. As we observed dysregulation of pathways associated with endocytosis in S305 mutant astrocytes, we tested whether increased *MAPT* Exon 10 inclusion may influence the internalization of tau. We therefore incubated S305 mutation and isogenic control astrocytes with 6 different isoforms of monomeric recombinant tau, labeled with pHrodo Red, and tracked tau internalization into cells over the course of 48 hours (Figure 7A-F, Figure S5A-C).

**Figure 7.**
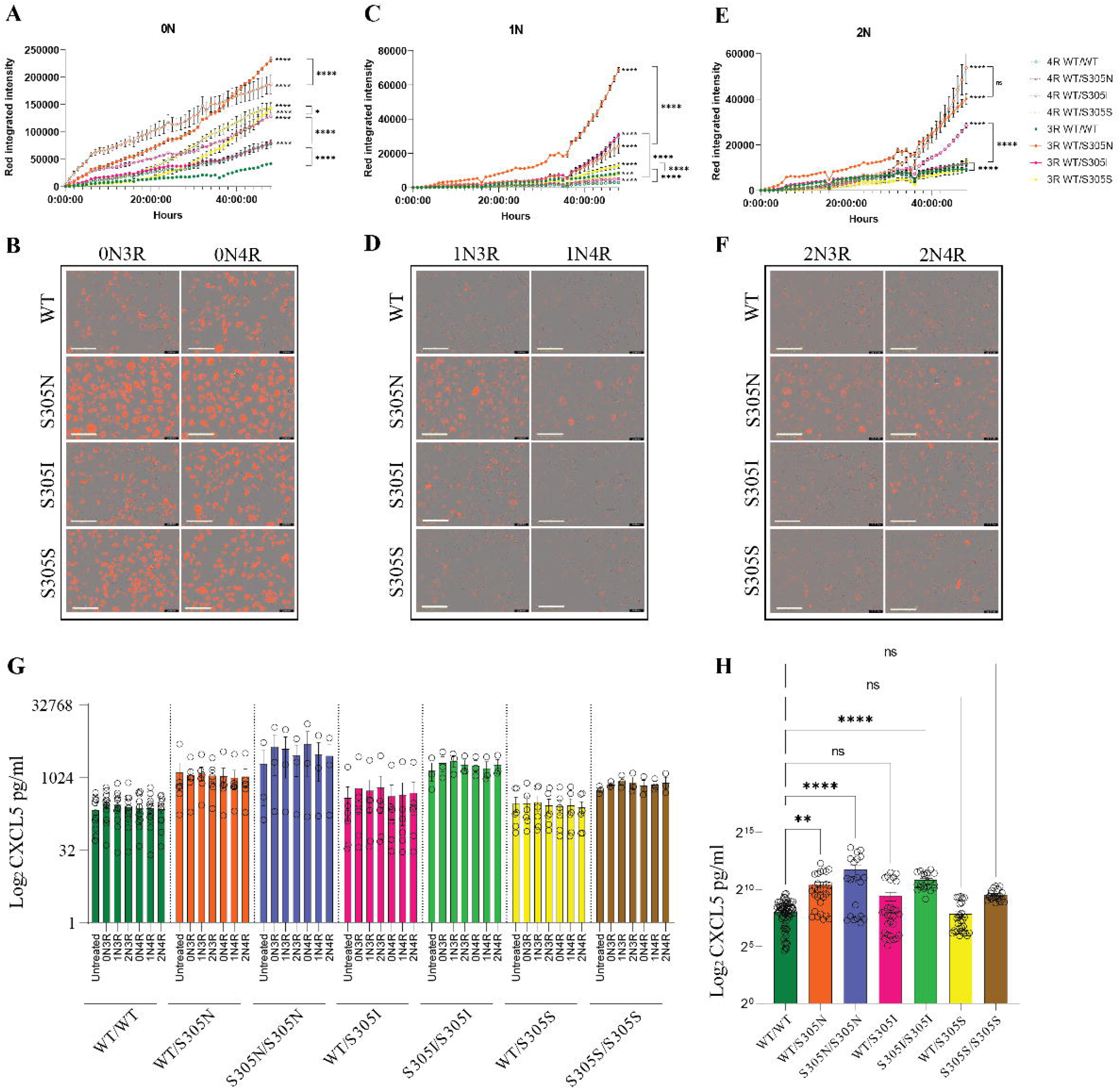
Astrocytes with *MAPT* S305 mutations are in an inflammatory state and rapidly internalize exogenous tau. **A.** Representative trace of integrated fluorescence intensity of pHrodo Red signal in S305 mutation astrocytes and isogenic controls treated with labelled 0N3R and 0N4R tau isoforms over 48 hours. Error bars = SEM. N = 4, each point is an average of 3 fields of view per well. Linear regression. **B.** Representative Incucyte images of S305 mutation astrocytes and controls at 48 hours following treatment with pHrodo-labelled 0N tau, quantified in ***A.*** Scale bar = 200µm. **C.** Representative trace of integrated fluorescence intensity of pHrodo Red signal in S305 mutation astrocytes and isogenic controls treated with labelled 1N3R and 1N4R tau isoforms over 48 hours. Error bars = SEM. N = 4, each point is an average of 3 fields of view per well. Linear regression. **D.** Representative Incucyte images of S305 mutation astrocytes and controls at 48 hours following treatment with pHrodo-labelled 1N tau, quantified in ***C***. Scale bar = 200µm. **E.** Representative trace of integrated fluorescence intensity of pHrodo Red signal in S305 mutation astrocytes and isogenic controls treated with labelled 2N3R and 2N4R tau isoforms over 48 hours. Error bars = SEM. N = 4, each point is an average of 3 fields of view per well. Linear regression. **F.** Representative Incucyte images of S305 mutation astrocytes and controls at 48 hours following treatment with pHrodo-labelled 2N tau, quantified in ***E***. Scale bar = 200µm. **G.** Log_2_ CXCL5 concentration (pg/ml) in iPSC-astrocyte culture media, corrected for cellular density, either untreated or following treatment with exogenous tau isoforms. Error bars = SEM. N = 3-12, each point represents an independent experiment averaging 3 technical replicates. Two-way ANOVA with Dunnett’s multiple comparison tests. **H.** Log_2_ CXCL5 concentration (pg/ml) in iPSC-astrocyte culture media, corrected for cellular density, pooled across all tau isoform treatment conditions for each genotype. Error bars = SEM. N = 21-70. One-way ANOVA with Dunnett’s multiple comparison tests.

We observed a significant effect of both genotype and tau isoform on the rate of astrocytic tau internalization (Figure 7A-F, Figure S5A-C) for all S305 mutations. Specifically, 0N isoforms were internalized faster than 1N isoforms, while 2N isoform internalization was inefficient across most genotypes (Figure 7A-F, Figure S5A-C). The preferential and specific uptake of 0N and 1N tau was consistent with our previous report that 2N tau was not observed in astrocyte pathologies in either PSP or AD brain^47^. The relationship between 3R and 4R isoforms was more complex; there was no consistent effect of 3R/4R tau for 0N or 2N isoforms across all three parent lines (Figures 7A-B, E-F, S5A, C). However, 1N3R was consistently internalized at a significantly faster rate than 1N4R tau across all mutations and parent lines (Figure 7C-D, Figure S5B), indicating both N and C-terminal splicing of *MAPT* have a combinatorial effect on its uptake. For all parent lines, S305 mutations showed significantly increased tau internalization compared to their isogenic control (Figure 7A-F, Figure S5A-C). Interestingly, comparison of each mutation on the same genetic background (F13505) revealed an association between the rate of tau internalization and the extent of *MAPT* exon 10 expression; tau uptake was fastest in S305N mutants, followed by S305I then S305S (Figure 7A-F), suggesting that 4R tau may directly influence astrocyte protein internalization..

As our gene expression data indicated a potential effect of *MAPT* S305 mutations on astrocytic basal inflammatory state, and inflammation has been described in PSP brain^48, 49^, we hypothesized that exogenous tau exposure could induce an inflammatory response in astrocytes that is modulated by S305 mutations. We therefore tested this hypothesis by conducting ELISAs for CXCL5 on the supernatant from tau-treated astrocytes (Figure 7G-H). Contrary to expectations, we did not observe any effect of tau treatment on the release of CXCL5 for any mutation (Figure 7G), indicating that exogenous tau exposure did not induce an inflammatory response. However, we did observe significantly increased CXCL5 at baseline in S305N and S305I mutant astrocytes (Figure 7H), indicative of an altered inflammatory state in these cells that could influence downstream function.

## DISCUSSION

Given the importance of 4R tau in both healthy and diseased brain, the very low levels of 4R tau expression in iPSC cultures have been a concern for complete and accurate modeling of tauopathy. These concerns have been partially addressed by utilizing iPSC lines carrying *MAPT* mutations in constitutively expressed exons, which are therefore present in the dominantly expressed 3R isoform^6, 8, 11, 14, 15, 30^. Additionally, iPSC-neurons carrying the *MAPT* P301L mutation, which is in Exon 10, still display tauopathy-relevant phenotypes despite very low 4R tau expression^7, 11^, indicating that Exon 10 inclusion, even at low levels, has a substantial impact on cellular function. Consistent with this assertion, 4R tau has been shown to more strongly stabilize microtubules compared to 3R, reduce cellular size, influence phosphorylation and modulate inflammatory responses^50–54^. It is therefore important to be able to model and manipulate the functional effects of 4R tau in a human neural cellular context in order to understand its contribution to pathogenic mechanisms across different cell types, and to facilitate screening of novel therapeutics in a more representative model of the human cellular environment.

For this purpose, we have developed a novel isogenic panel of iPSC lines, derived from four different donors, including heterozygote and homozygote *MAPT* S305N, S305I and S305S splicing mutations and paired isogenic corrected controls, totaling 31 clones. We found that the S305N mutation had the most severe effect on Exon 10 inclusion, with up to 70% 4R transcripts from as early as 4 weeks of neuronal differentiation. Significantly increased 4R tau was also present at the protein level, with almost exclusively 4R expression in homozygous S305N mutants by 8 weeks of differentiation. This severity is likely due to the effect that the S305N mutation has on the stability of the Exon 10 – Intron 10 hairpin structure, as it has previously been identified as one of the most destabilizing modifications possible in this region^28^. S305I mutants were second in the proportion of Exon 10 inclusion; however, there was substantial variability in heterozygous mutants compared to the homozygotes. This was possibly a result of donor line variability, as the healthy control iPSC line (F13505) into which we introduced the S305I mutation resulted in less Exon 10 inclusion compared to the clones derived from a mutation carrier patient (75.11). This suggests that interaction with individual genetic backgrounds may also influence the severity of S305I splice mutations, as has been suggested for other pathogenic mutations^55, 56^. Regardless, we were able to observe consistent transcriptomic and functional effects of this mutation across donor lines, despite variability in 4R tau expression.

The synonymous S305S mutation had the least severe and minimal influence on Exon 10 inclusion, although this increase was sufficient to induce functional and transcriptomic changes comparable to S305N and S305I mutation lines. This reduced severity may be reflective of the clinical syndrome, in which the age of onset in affected carriers was into the 40s and 50s^26, 27^, compared with a younger age of onset in their 30s and subsequent death within 2 years for currently reported S305N and S305I carriers^24, 25^. The milder effect of S305S parallels the characterization of *MAPT* IVS10+16 mutations, which destabilize the Exon 10 – Intron 10 hairpin loop without introducing an amino acid change, and have a similar age of onset around 40-50 years old^57^. However, unlike IVS10+16 mutation iPSC-neurons, which show progressive Exon 10 inclusion over several months of differentiation^19, 20^, S305S, as well as S305N and S305I mutations appeared to have an early and sustained influence on the proportion of Exon 10 splicing. As we cultured our neurons for only 8 weeks, longer maintenance would be required to confirm no additional changes in the proportion of 4R tau expression over a prolonged period across all three S305 mutations.

This early aberrant expression of 4R tau has implications for how we may consider susceptibility for neurodegenerative disorders, and the developmental impact of pathogenic mutations. It has been hypothesized that the preference for 3R tau expression in human fetal brain is due to its reduced ability to stabilize microtubules, and is therefore more permissive of the synaptic plasticity and remodeling that is necessary for healthy brain development^4, 58^. Indeed, overexpression of 4R tau in SH-SY5Y cells was found to promote neuronal maturation compared to 3R tau expression that did not^59^. We may therefore expect that expression of 4R tau too early in neuronal development will have fundamental effects on cellular function with downstream consequences for disease susceptibility later in life. This potential mechanism may be most apparent in S305S neurons, as there remained significant transcriptomic changes associated with the mutation at 8 weeks of differentiation, even though the proportion of Exon 10 inclusion in isogenic corrected controls was comparable to the mutants by this time point.

Curiously, some of the most prominently affected pathways across all three mutations converged on downregulation of *Neurotransmitter* and *Glutamate Signaling*. This came as a surprise, as we and others have observed upregulation of equivalent pathways in iPSC models with different *MAPT* mutations, as well as increased neuronal firing and hyperexcitability^6, 7, 17, 60^. However, further inspection of these genes suggested downregulation of inhibitory GABAergic signaling pathways, as well as enrichment for pathways associated with hyperexcitability such as seizure susceptibility. We observed early neuronal activity and synaptic maturation in S305 mutants compared to their isogenic controls, consistent with observations from V337M, N279K and P301L *MAPT* mutation iPSC-neurons^7, 60^. A recent electrophysiological study of IVS10+16 mutation neurons suggested increased 4R tau caused impaired neuronal excitability^17^, although these studies were conducted much later in differentiation than in our experiments (∼200 days compared to 3-8 weeks). This discrepancy may therefore be indicative of a progressive phenotype that switches from hyperexcitability to impairment with age, as has been previously suggested^6, 7^. The reciprocal relationship we observed between downregulation of neurotransmitters and synaptic firing and upregulation of seizure-related pathways, coupled with increased electrophysiological activity in S305 mutants, suggests that early neuronal hyperexcitability commonly observed in different *MAPT* mutation models may occur via different mechanisms, ultimately converging on alteration of Excitatory/Inhibitory balance. Similar to observations in mouse brain^61, 62^, we observe increased localization of tau and phosphorylated tau to synaptosomes in S305 mutant neurons, where it has the potential to alter or disrupt synaptic function via modulation of presynaptic vesicle recycling, endocytosis impairment, and by post-synaptic interaction with Fyn^63–65^.

Although the S305N mutation lines tended to show the most extreme phenotypes and largest gene expression changes compared to S305I and S305S, it is unclear how much of this effect could be attributed to the particular amino acid change, or to their substantially higher proportion of 4R tau expression. However, the divergent mechanistic changes between missense (S305N/S305I) and synonymous (S305S) mutations may provide an indication of WT 4R vs mutant 4R effects. One of the most apparent differences consistently observed between missense and synonymous mutations was in mitochondrial bioenergetics. Recently, WT tau (2N4R) was demonstrated to directly interact with different components of the mitochondrial membrane, but specific protein interactions changed with the introduction of P301L or V337M *MAPT* mutations, such that they altered mitochondrial function^32^. It is therefore plausible that the S305N and S305I amino acid changes modify the tau mitochondrial interactome, resulting in divergent functional effects on bioenergetics compared to a WT 4R protein. Our data from the S305S mutation neurons suggests that there are likely to be additional differences between 3R and 4R tau in their interaction with the mitochondrial membrane, although the specific effect of isoform has not yet been explored. However, general suppression of mitochondrial bioenergetics in these cells may indicate an inhibition of mitochondrial function caused by 4R tau, rendering neurons less able to cope with alterations in energetic demands.

Notably, the most significant effect of missense S305 mutations was on non-mitochondrial respiration, which was increased in both heterozygous and homozygous mutants, and could reflect the use of glycolysis as an energy source rather than, or in addition to, mitochondrial oxidative phosphorylation. This is a curious observation, as the switch from glycolysis in neural progenitor cells to oxidative phosphorylation in neurons has been well documented^32, 66^, and would suggest that energy production in S305N and S305I mutant neurons is not efficiently undergoing this switch. Alternatively, an increase in glycolysis may reflect a response to an inflammatory state^67^ in order to deal with increased energetic demand of cellular stress. This increase in non-mitochondrial respiration may explain the susceptibility to oxidative stress and excitotoxicity observed in other *MAPT* mutation models^6, 8^, as neurons are unable to fully utilize oxidative phosphorylation to cope with additional energy demands, and are already primed and susceptible to stress. However, a trend towards increased basal and maximal respiration, and significant enrichment of oxidative phosphorylation gene expression pathways in these cells is not fully consistent with this hypothesis, but could reflect a compensatory response to inefficient ATP production, which was unchanged in these neurons.

Consistent with the assertion that missense S305 mutations may be inducing an inflammatory state in neurons, whereas the synonymous S305S does not, we observed a similar pattern in S305 mutation astrocytes. Both S305N and S305I mutations were associated with increased basal CXCL5, a cytokine associated with aged astrocytes, and the promotion of neutrophil chemotaxis in response to inflammatory stimuli^68, 69^. Increased CXCL5 detection in postmortem cerebrospinal fluid (CSF) has recently been observed in cases of chronic traumatic encephalopathy (CTE)^70^ and AD-related mild cognitive impairment (AD-MCI)^71^, supporting the relevance of this cytokine to tauopathy pathophysiology, although the mechanism by which astrocytic endogenous tau may promote CXCL5 release is unclear.

Recently, more attention has been given to the role of astrocytes in tauopathy, as well as the recognition that even very low levels of *MAPT* expression in these cells has the potential for cell autonomous and non-cell autonomous effects^41, 72, 73^. Consistent with recent studies in IVS10+16 iPSC-astrocytes^41^, we also observe downregulation of pathways associated with glutamate and neurotransmitter signaling, consistent with the hypothesis that *MAPT* mutation astrocytic dysfunction contributes to neuronal excitotoxicity by being less able to clear excess glutamate away from the synaptic cleft^74^. Others have also noted that iPSC-astrocytes splice *MAPT* differently than neurons, and the effect of the IVS10+16 mutation on Exon 10 inclusion is exacerbated in astrocytes^73^. We observe the same phenomenon in S305 mutation astrocytes, although our ISOseq data also indicates that *MAPT* splicing regulation is generally different in astrocytes compared to neurons across the full length of the gene, not just at Exon 10. Specifically, we observe a wider diversity of transcripts, and the expression of longer transcripts in astrocytes compared to neurons, with increased inclusion of Exons 2, 3 and 10, but also Exons 6 and 4a, which are not typically considered to be expressed in human brain. Exon 4a expression is particularly interesting, as its expression is considered to be restricted to the peripheral nervous system (PNS) and projecting neurons, and not the central nervous system^44, 75^. However, low level astrocyte-specific expression may have precluded its detection to date. Very little is known about the function of Exon 4a, although it has been hypothesized to provide additional stabilization to microtubules that facilitates longer-range axonal projections required in the PNS^44^. A potential function for Exon 4a inclusion in astrocytes is unclear. Given the variability in our isoform data, it was not possible to determine whether S305 mutations alter the splicing of other *MAPT* exons in astrocytes. It remains to be seen what the functional impact of astrocytic *MAPT* splicing may be, what the effect of differential isoform usage is in these cells, and the relationship to disease pathogenesis. It should also be noted that we have made these observations only in iPSC-derived astrocytes, therefore confirmation of altered *MAPT* splicing in human brain-derived astroglia is required to confidently conclude that astrocytes process and splice *MAPT* in this manner.

Finally, as tau accumulation in astrocytes is a characteristic neuropathological feature of several primary tauopathies, and our gene expression data implicated alterations in endocytic function, we decided to explore whether *MAPT* S305 mutations influenced the astrocytic internalization of exogenous tau protein. Astrocytic internalization of monomeric tau has been shown before in rodent astrocytes^76^, but we now demonstrate in a human system that *MAPT* mutations influence uptake of tau in a manner that correlates with the proportion of Exon 10 expression, i.e. S305N internalization was fastest, followed by S305I, then S305S and finally by isogenic corrected controls, therefore implicating 4R tau in astrocyte protein internalization capabilities. Surprisingly, we saw a significant effect of tau isoform on the rate of astrocytic internalization, with 0N isoforms most rapidly being processed, and 2N isoforms largely remaining un-phagocytosed. This pattern of internalization is consistent with what has been observed in human brain pathology: tau pathology in AD and PSP brain has been shown to primarily consist of 0N and 1N isoforms, whereas 2N isoforms were not present within glia^47^. These differences may therefore reflect the differential ability for exogenous N-terminal isoforms to enter cells and subsequently accumulate. We did not observe an inflammatory response to treatment with exogenous tau protein, however this may be due to our use of monomeric recombinant tau that did not have any posttranslational modifications. It therefore remains to be seen how S305 mutation astrocytes may respond to oligomeric or hyperphosphorylated protein. Alternatively, as an inflammatory state can promote phagocytosis^77^, and S305 mutation lines with increased tau internalization had increased CXCL5 release at baseline, it is possible that inflammation occurs prior to, and subsequently exacerbates tau internalization and accumulation in astrocytes. Indeed, glial inflammation has been found to exacerbate tau pathology in mouse models^78, 79^. Additionally, as protein sequence and length influences astrocytic uptake of different tau isoforms, it therefore remains to be seen whether S305 mutations differentially influence the internalization of mutant protein, or other substrates that could affect downstream cellular function.

In conclusion, we have developed a unique and valuable panel of isogenic *MAPT* S305 mutation iPSC lines, which express unprecedented levels of 4R tau very early in iPSC-derived neurons and astrocytes. These lines recapitulate previously observed neuronal phenotypes such as hyperexcitability and alterations in synaptic function. They also highlight functional differences between missense *MAPT* mutations and pathological WT 4R tau expression in pathways such as mitochondrial bioenergetics, which may have implications for how we understand sporadic compared to familial disease pathogenesis. We also highlight the functional importance of *MAPT* expression in astrocytes, and implicate 4R tau in the regulation of inflammatory and phagocytic processes. We believe that these lines will be valuable to tauopathy researchers enabling a more complete understanding of the pathogenic mechanisms underlying 4R tauopathies across different cell types.

## METHODS

### iPSC Reprogramming

WT/S305N and WT/S305I peripheral blood mononuclear cells were identified as part of the ALLFTD study (https://www.allftd.org/), and were requested and obtained from the National Centralized Repository for Alzheimer’s Disease (NCRAD; https://ncrad.iu.edu/). WT/S305S lymphoblastoid cell lines (LCLs) were obtained from the Sydney Brain Bank (https://sbb.neura.edu.au/). PBMCs and LCLs were reprogrammed to iPSCs by the Icahn School of Medicine at Mount Sinai Stem Cell core facility, using Sendai virus mediated gene transfer of *OCT4*, *SOX2*, *KLF4* and c-*MYC*. Following reprogramming, karyotypic normality of three parent clones was confirmed by G-band karyotyping conducted by WiCell (www.wicell.org), and mutation status was confirmed by Sanger sequencing (see below for details). Expression of pluripotency markers was assessed by immunofluorescence (see below for details).

### CRISPR/Cas9 genome editing

Human iPSCs were edited using the pSpCas9(BB)-2A-GFP (PX458) construct backbone (Addgene #48138) described in Ran *et al* 2013^80^. For WT correction of S305 mutation lines, specific guide RNAs (gRNAs) were designed against the mutant allele using the Zhang lab design tool (crispr.mit.edu), and guides with the highest quality scores and more than 3bp mismatches with other genomic loci were selected. For the generation of homozygous mutants and introduction of S305 mutations onto a WT control background, guides were designed against the WT allele with the same method. Guides were cloned into the CRISPR/Cas9 vector backbone using the Zhang lab protocol (addgene.org/crispr/zhang) and correct insertion was confirmed by Sanger sequencing using primers for the human U6 promoter. Validation of correct targeting and cutting activity was validated in HEK293T cells using the Surveyor nuclease assay (IDT) before use in iPSCs. Repair templates (single stranded oligodeoxynucleotides (ssODN)) were designed to span 70 nucleotides either side of the Cas9 cleavage site. ssODNs were designed with a synonymous protospacer adjacent motif (PAM) site modification to reduce the chance of multiple cutting events (V300V; GUC > GUA, Figure S6A). Using computational modeling with experimental chemical and microarray mapping constraints, Lisowiec *et al*.^81^ proposed a 14-nt hairpin and a 25-nt hairpin containing the 5’ splice site at the Exon 10-Intron 10 junction (Figure S6B). The V300V mutation was predicted to stabilize this structure (Figure S6B). This was confirmed by introduction of the mutation into a *MAPT* Exon 10 minigene^82^ by site directed mutagenesis and expression in SH-SY5Y cells (Figure S6C). S305S, S305N and S305I mutations all substantially increased Exon 10 inclusion in this system (Figure S6C). The PAM site edit was not incorporated in all clones, therefore summary of PAM edit status is included in Table 1. All guide RNA sequences are provided in Table S1.

For editing, iPSCs were maintained in DMEM/F12 with 10% FBS and 10µM Y27632, without antibiotics, overnight. Cells were then dissociated into a single cell suspension in Accutase (Innnovative Cell Tech), and 2x10^6^ cells were electroporated with the Neon transfection system (ThermoFisher Scientific) in 100µl tips with 4µg gRNA-Cas9 vector and 40µM ssODN. Cells were electroporated at 1400V for 20ms, with 2 pulses. Electroporated cells were seeded into a 10cm dish in DMEM/F12, 10% FBS and 10µM Y27632 for recovery. After 24 hours, media was replaced daily with StemFlex (ThermoFisher Scientific) containing gradually reducing concentrations of Y27632. When recovered, 288 individual clones (3 x 96w plates) were selected using Collagenase (ThermoFisher Scientific) dissociation and manual selection under a microscope. Clones were seeded into 96-well plates for recovery and expansion. Each plate was then split 1:2 using ReLeSR (StemCell Technologies), with half of the cells used for DNA extraction for Sanger sequencing, and the other half cryopreserved until mutation status could be confirmed.

### Sanger Sequencing

DNA was extracted from 96-well plates of cells using QuickExtract (Epicentre) by incubation at 65°C for 6 minutes, followed by 2 minutes at 98°C. The region spanning the targeted mutation was amplified using Phusion mastermix (Cell Signaling Technology) with the following primers: Forward 5’-GACTCAACCTCCCGTCACTC -3’, Reverse 5’-GGGACACCCCCTCCTAGAAT -3’. PCR products were confirmed by Tapestation (Agilent), and cleaned up with ExoSAP-IT (ThermoFisher Scientific). Forward and reverse sequencing reactions were carried out using 20ng of PCR product and BigDye Terminator v3.1, with the same primers used for PCR amplification. Before sequencing, products were cleaned up using the BigDye Xterminator purification kit (ThermoFisher Scientific) and were assessed using the SeqStudio Genetic Analyzer (ThermoFisher Scientific). Sequences were visualized using Chromas (https://technelysium.com.au/wp/chromas/).

### iPSC culture and differentiation

iPSCs were maintained on Matrigel (Corning)-coated plates in StemFlex (ThermoFisher Scientific) supplemented with 1% penicillin/streptomycin (ThermoFisher Scientific), and were passaged with ReLeSR (StemCell Technologies). iPSCs were differentiated to neural progenitor cells (NPCs) and were MACS sorted to deplete neural crest cells, as previously described^83^. For neuronal differentiation, NPCs were incubated with BrainPhys (StemCell Technologies) supplemented with 1xB27 without vitamin A, 1xN2 (both ThermoFisher Scientific), 20ng/ml BDNF, 20ng/ml GDNF (both R&D), 200µM ascorbic acid, 250µg/ml cyclic AMP (both Sigma Aldrich) and 1% penicillin/streptomycin. Neurons were maintained for up to 8 weeks, with media changes every other day. For astrocyte differentiation, NPCs were seeded sparsely into Matrigel-coated plates and treated with Astrocyte media (ScienCell) as previously described^84^. Astrocytes were collected or used for experiments at 30 days of differentiation.

### Immunofluorescence

Cells were fixed with 10% Formalin (Sigma Aldrich) for 15 minutes at room temperature, then washed three times with PBS. They were then incubated with 0.1% Triton X-100 in PBS for 30 minutes with gentle rocking, followed by three additional PBS washes, and incubation with 1% bovine serum albumin (BSA) in PBS for a minimum of 30 minutes at room temperature. Cells were then incubated with primary antibodies prepared in 1% BSA in PBS overnight at 4°C, followed by three PBS washes and incubation with the appropriate secondary antibodies at a 1:100 dilution in 1% BSA in PBS for a minimum of 2 hours at room temperature. Secondary antibodies were then removed, and cells were incubated for 10 minutes at room temperature with 300nM DAPI prepared in PBS. Three PBS washes were repeated, and cells were stored in PBS at 4°C until ready for imaging. For pluripotency confirmation, the following antibodies were used: TRA-1-81 (Cell Signaling Technologies 4745, 1:1000), SSEA4 (Cell Signaling Technologies 4755, 1:1000), TRA-1-60 (Cell Signaling Technologies 4746, 1:1000), Nanog (Cell Signaling Technologies D73G4, 1:400), SOX2 (Cell Signaling Technologies 3579S, 1:400) and OCT4A (Cell Signaling Technologies 2840S, 1:400). Secondary antibodies were AlexaFluors Goat-anti-mouse 488 and Donkey-anti-Rabbit 568, all used at 1:100 and purchased from ThermoFisher Scientific.

### Live lysosome labelling

Lysotracker (ThermoFisher Scientific) was added to astrocyte culture media at a final concentration of 100nM and was incubated for 30 minutes at 37°C. Lysotracker-media was then removed and replaced with fresh astrocyte culture media containing 1µM Lysosensor (ThermoFisher Scientific). Cells were incubated for 1 minute at 37°C, then washed three times with fresh astrocyte media and immediately imaged on a Leica DMIL LED Inverted Routine Fluorescence Microscope with a 20x objective. Lysotracker and Lysosensor images underwent automated integrated density analysis with a set minimum grey intensity threshold for positive signal across all images, using ImageJ v1.53t.

### SDS-PAGE gel electrophoresis and western blot

For whole protein lysate analysis, cells were suspended in Cell Lysis Buffer (Cell Signaling Technology) supplemented with 10µM PMSF and 1x protease/phosphatase cocktail (ThermoFisher Scientific) and sonicated on ice, followed by centrifugation at 13,000 x g for 10 minutes at 4°C to pellet debris. Protein concentration was determined by BCA assay (ThermoFisher Scientific).

For synaptosome and cytosolic fractions, cells were lysed in Syn-Per Synaptic Protein Extraction reagent (ThermoFisher Scientific) and centrifuged first at 1,200 x g for 10 minutes at 4°C to remove debris, then a further 15,000xg for 20 minutes at 4°C to pellet synaptic proteins. The pellet was resuspended in PBS and retained as the synaptosome fraction, while the remaining supernatant was the cytosolic fraction. Protein concentrations were determined by BCA assay.

10-20µg of protein per sample was incubated with 1x Reducing Agent and 1x LDS sample buffer (both ThermoFisher Scientific) at 70°C for 10 minutes, before being immediately loaded onto BOLT 4-16% Bis-Tris gels in 1x MES buffer (ThermoFisher Scientific). Electrophoresis was carried out for 25 minutes at 200V before blotting onto nitrocellulose membranes using the iBlot system (ThermoFisher Scientific). Membranes were blocked for a minimum of 30 minutes at room temperature in 5% milk in PBS-T. Primary antibodies were prepared at the following dilutions in 5% milk in PBS-T: Da9 (total tau) 1:500, PHF1 (p-tau_S396_) 1:500 (both kind gifts from Dr. Peter Davies), Synapsin1 1:1000 (Synaptic Systems 106011), Homer1 1:000 (Synaptic Systems 160003), GAPDH 1:10,000 (Abcam ab181602), and were incubated with the membrane overnight at 4°C with rocking. Membranes were washed 3x with PBS-T. Membranes were then incubated with either HRP Goat Anti-Rabbit or HRP Horse Anti-Mouse secondary antibodies (Vector laboratories) at a dilution of 1:20,000 in 5% milk in PBS-T for one hour at room temperature with gentle rocking. Following three additional washes, membranes were then incubated with WesternBright ECL HRP substrate (Advansta) for 3 minutes before imaging on a UVP ChemiDoc. For re-staining, blots were incubated in Restore PLUS stripping buffer (ThermoFisher Scientific) for 15 minutes at room temperature, followed by one wash in PBS and re-blocking.

### qRTPCR

Cell pellets were collected in cold PBS by scraping, and RNA was extracted using the Qiagen RNeasy mini RNA extraction kit, then reverse transcribed using the high capacity RNA-to-cDNA kit (ThermoFisher Scientific). For qRTPCR of 3R and 4R *MAPT* transcripts, amplification was carried out using SybrGreen mastermix (ThermoFisher Scientific) with the following primers: *MAPT* 4R Forward 5’-CGGGAAGGTGCAGATAATTAA-3’, Reverse 5’-GCCACCTCCTGGTTTATGATG-3’; *MAPT* 3R Forward 5’-AGGCGGGAAGGTGCAAATA-3’, Reverse 5’-GCCACCTCCTGGTTTATGATG-3’.

### RNA-seq

RNA was isolated as described above, and library preparation with poly-A selection followed by sequencing with 150 base pair paired-end reads was carried out at Genewiz. Sequenced reads were trimmed for Illumina TruSeq adapters, and aligned to the human hg38 genome using the STAR aligner^85^ for quantification of gene-level read counts. Genes were filtered by expression abundance for counts per million (CPM) of > 0.1 in ≥ 5 samples. Library sizes were estimated by the trimmed mean of M-values normalization (TMM) method^86^ using the R edgR package v3.26.8. Differential gene expression was predicted by linear mixed model analysis using the variancePartition package v1.14.1^87^ in order to account for sample correlation in isogenic lines. False discovery rate (FDR) of the differential expression test was estimated using the Benjamini-Hochberg (BH) method^88^. Gene set enrichment analysis (GSEA) was performed using the Molecular Signatures Database (MSigDB) Gene Ontology (GO) annotations of canonical pathways, biological processes, cellular compartments and molecular function, with 10,000 permutations. Additional pathway analyses were carried out using Ingenuity Pathway Analysis (Qiagen Inc.; https://digitalinsights.qiagen.com/products/ingenuitypathway-analysis). Exon 10 PSI values were calculated from aligned BAM files using Mixture of Isoforms (MISO) software^89^ with default parameters.

### *MAPT* targeted ISOseq

RNA was submitted to the Icahn School of Medicine at Mount Sinai Genomics CoRE for single molecule real time (SMRT) isoform sequencing (ISOseq) on the PacBio RS II platform using the following primers: Forward 5’-ATG GAA GAT CAC GCT GGG AC-3’, Reverse 5’-GAG GCA GAC ACC TCG TCA G-3’. Raw sequencing reads were passed through the ISOseq3 pipeline to detect full-length transcripts expressed in each sample. Beginning with raw subreads, single consensus sequences were generated for each *MAPT* amplicon with a SMRT adapter on both ends of the molecule. SMRT adapter sequences were then removed and *MAPT*-specific primer sequences were identified to orient the isoforms. Isoforms were subsequently trimmed of poly(A) tails and concatemers were identified and removed. Isoform consensus sequences were then predicted using a hierarchical alignment and iterative cluster merging algorithm to align incomplete reads to longer sequences. Finally, clustered isoform sequences were polished using the arrow model and binned into groups of isoforms with predicted accuracy of either ≥ 0.99 (high quality) or < 0.99 (low quality). The resulting isoforms were aligned to hg38 using the GMAP aligner^90^.

### MEA

NPCs were seeded at a density of 25,000 cells per well in 48-well CytoView plates (Axion Biosystems) and differentiated into neurons as described above. Every week between 2- and 8-weeks of differentiation, spontaneous activity was measured with a 10 minute recording on the Axion Maestro using default neural real time spontaneous activity settings, and data was processed using the Axion AxiS Software with default Neural activity settings.

### Seahorse assays

NPCs were seeded at a density of 50,000 cells per well into Seahorse XF96 96-well plates (Agilent), and were differentiated into neurons as described above. Mitochondrial bioenergetics were assessed at 6 weeks of differentiation using the Mitochondrial Stress Test measured on a Seahorse XFe96 Analyzer (both Agilent). Stress tests were carried out according to manufacturer’s instructions and cycling parameters. Briefly, neuronal media was changed to XF DMEM (Agilent) containing 1mM pyruvate, 2mM glutamine and 10mM glucose. Cells were equilibrated to the new media for 1 hour at 37°C in the absence of CO_2_ prior to starting the assay. The plate underwent 3 mix/measurement cycles to establish baseline OCR, followed by treatment with 1.5µM oligomycin followed by an additional 3 cycles. Cells were then exposed to 0.5µM FCCP and measured over 3 cycles, then were finally treated with 0.5µM Rotenone/Antimycin A and OCR was measured over a final 3 cycles. After assay completion, culture media was removed and replaced with Methylene Blue and incubated for 1 hour at room temperature to fix and stain cells. Methylene Blue was removed and the plate was thoroughly washed with distilled water, before cells were lysed with 4% acetic acid in 40% methanol. The resulting de-stain was transferred to a new plate and absorbance at 595nm was measured using a Varioskan microplate reader (ThermoFisher). The resulting values were applied as a scaling factor in Seahorse Wave analyzer software (V2.6.3.5) to correct OCR measurements for cell density. All OCR calculations were generated by the Seahorse Wave software.

### Tau uptake assays

Monomeric recombinant tau of each of the 6 major isoforms was purchased from rPeptide, and was labelled using the pHrodo Red Microscale Protein Labeling kit (ThermoFisher Scientific). 100µg protein (1mg/ml) was incubated with 100µM sodium bicarbonate. 3µl of 2mM pHrodo iFL STP ester reactive dye was added to the sample, and incubated at room temperature for 15 minutes. Excess dye was removed from the sample by centrifugation for 5 minutes at 1,000 x g through a spin column containing gel resin

For the tau uptake assay, iPSC-astrocytes were seeded into 96-well plates at a density of 50,000 cells per well. After 24 hours, cells were then incubated with astrocyte media containing 1.5µg of the relevant pHrodo-labeled tau isoform or the equivalent volume of PBS. Treated plates were immediately placed into the IncuCyte live cell imaging system (EssenBio), and three fields of view per well were imaged every hour for 48 hours with a 20x objective. Image analysis was carried out in IncuCyte software for automated measurement of red integrated intensity and cell confluence.

### ELISAs

Following exogenous tau treatment and imaging, conditioned media was collected from iPSC-astrocytes for assessment of CXCL5 secretion by sandwich ELISA (Abcam ab212163) according to manufacturer’s instructions. Absorbance was measured on a Varioskan microplate reader (ThermoFisher Scientific) and normalized to cell confluence as measured by Incucyte imaging at the time of media collection. CXCL5 concentration was determined by comparison to a positive control standard curve.

## Statistical analysis

RNA-seq and immunofluorescence data were analyzed as described above. Western blot protein bands were quantified by densitometry analysis in ImageJ, and normalized to GAPDH for each sample. For ISOseq data, the expression of each isoform was calculated as a proportion of all detected isoforms in that sample, and average expression for each genotype was calculated. qRTPCR gene expression was analyzed using the ΔΔCt method, and expression was normalized to β-actin as endogenous controls, then a ratio between 4R and 3R expression was calculated for each sample. Comparisons across time and genotypes were carried out using the appropriate two-way ANOVA and post-hoc testing, and comparisons between genotypes at single time points were carried out using one-way ANOVA and post-hoc comparisons for multiple test correction. Incucyte time course data was analyzed using simple linear models. All analyses were carried out in either GraphPad Prism 9, or R v4.2.2. For cell culture experiments, tests were either conducted on all available clones, or on one clone per genotype per parent line with three independent rounds of differentiation, averaged from multiple technical replicates. Specific statistical tests and N for individual assays are described in the relevant figure legends.

## Supporting information

SFig1

SFig2

SFig3

SFig4

SFig5

SFig6

## ACKNOWLEDGEMENTS

We thank the research subjects and their families for their generous participation. Data collection and dissemination of the data presented in this manuscript was supported by the ALLFTD Consortium (U19: AG063911, funded by the National Institute on Aging and the National Institute of Neurological Diseases and Stroke) and the former ARTFL & LEFFTDS Consortia (ARTFL: U54 NS092089, funded by the National Institute of Neurological Diseases and Stroke and National Center for Advancing Translational Sciences; LEFFTDS: U01 AG045390, funded by the National Institute on Aging and the National Institute of Neurological Diseases and Stroke). The manuscript has been reviewed by the ALLFTD Executive Committee for scientific content. The authors acknowledge the invaluable contributions of the study participants and families as well as the assistance of the support staffs at each of the participating sites. We are also grateful to the Sydney Brain Bank for contributing S305S mutation lymphoblast cell lines (LCLs) to this study. LCLs were obtained from the New South Wales Brain Bank at Neuroscience Research Australia, which is supported by the University of New South Wales and Neuroscience Research Australia. We thank Dr. Celeste Karch for providing lines generated by the Washington University facility.

This work was supported by funding from the BrightFocus Foundation (KRB), Association for Frontotemporal Degeneration (KRB), CurePSP (KRB), and the Rainwater Charitable Foundation (AMG, KRB).

We thank the Genetics and Stem Cell CoREs at the Icahn School of Medicine at Mount Sinai for iPSC reprogramming and conducting targeted ISOseq data analysis. We are also very grateful to Drs. Jonathan Chen and Matt Disney (Scripps, Florida) for their assistance in calculating hairpin loop free energies and input regarding RNA stability.

## AUTHOR CONTRIBUTIONS

Conceptualization: KRB, AMG

Methodology: KRB, CP, DAP, SAW, LO, YL, JLC

Validation: KRB, CP, DAP, SAW, LO

Formal Analysis: KRB, CP, YL, JLC

Investigation: KRB, CP, DAP

Resources: KRB, AMG, MDD

Data Curation: KRB, CP, YL

Writing – Original draft: KRB

Writing – Review and editing: KRB, CP, DAP, SAW, LO, YL, AMG, JLC, MDD

Visualization: KRB, JLC

Supervision: AMG

Funding Acquisition: KRB, AMG

## SUPPLEMENTARY INFORMATION

### Supplementary Figure Legends

**Figure S1. Karyotyping and sequencing of CRISPR-edited clones (related to Figure 1)**

G-band karyotyping and Sanger sequencing traces from all CRISPR-edited and non-edited clonal controls generated from the four parent iPSC lines.

**Figure S2. *MAPT* S305 mutations have both shared and divergent effects on Exon 10 inclusion and differentially expressed gene pathway enrichment (related to Figures 2 and 3).**

**A.** *MAPT* expression (transcripts per million, TPM) from RNA-seq data of 4, 6 and 8 week old iPSC-neurons per clone, separated by mutation. Error bars = SEM. N = 2-8. Two-way mixed model ANOVA, correcting for Parent Line, with Dunnett’s post-hoc test correction.

**B.** 4R:3R ratio calculated from qRTPCR analysis of Exon 10+ and Exon 10-transcripts across all clones at each time point. Error bars = SEM. N = 2-8. Two-way ANOVA. *p < 0.05, **p < 0.01, ***p < 0.001, ****p < 0.0001.

**C-D.** Quantification of ***C.*** total tau and ***D.*** p-tau (S396), normalized to GAPDH across all mutations and time points, as represented in Figure 2C. Error bars = SEM. Two-way ANOVA with Bonferroni post-hoc testing compared to isogenic controls

**E-F.** Overlap of the number of significantly enriched pathways from GSEA analysis between S305 ***E*.** heterozygous and ***F.*** homozygous mutations at each time point, compared to isogenic controls.

**Figure S3. Neuronal synaptic maturity is accelerated in *MAPT* S305 mutation lines (related to Figure 4)**

**A.** Average number of spikes derived from multi-electrode array analysis of isogenic sets of S305N, S305I and S305S iPSC-neurons every week from 3-8 weeks of age. Error bars = SEM. N = 3 independent differentiations, averaged from 3 technical replicates per genotype/clone. Two-way ANOVA.

**B.** Quantification of western blots as represented in ***C,*** for total tau, p-tau (S396), Synapsin1 and Homer1, normalized to GAPDH loading control in 6-week old S305I mutation cytosol (C) and synaptosome (S) fractions compared to isogenic controls. Error bars = SEM. N = 3. One way ANOVA with Bonferroni post-hoc testing.

**C.** Representative western blot of cytosolic and synaptosome fractions derived from 6-week old S305I mutation neurons and isogenic controls, labelled for total tau and phospho-tau, as well as synaptic markers Synapsin1 and Homer1. GAPDH used as loading control.

**D.** Quantification of western blots as represented in ***E,*** for total tau, p-tau (S396), Synapsin1 and Homer 1, normalized to GAPDH loading control in 6-week old S305S mutation cytosol (C) and synaptosome (S) fractions compared to isogenic controls. Error bars = SEM. N = 3. One way ANOVA with Bonferroni post-hoc testing. *p < 0.05, **p < 0.01, ***p < 0.001

**E.** Representative western blot of cytosolic and synaptosome fractions derived from 6-week old S305S mutation neurons and isogenic controls, labelled for total tau and phospho-tau, as well as synaptic markers Synapsin1 and Homer1. GAPDH used as loading control.

**Figure S4. *MAPT* S305 neurons display altered mitochondrial phenotypes (related to Figure 5).**

**A-B.** Quantification of mitochondrial ***A.*** spare respiratory capacity and ***B.*** proton leak relative to isogenic controls in *MAPT* S305 mutation neurons at 6 weeks. Error bars = SEM. N = 3 independent differentiations, averaged from 3 technical replicates each time. One-way ANOVA with Fisher’s LSD posthoc testing. **p < 0.01

**Figure S5. iPSC-astrocytes with *MAPT* S305 mutations have altered tau internalization (related to Figure 7).**

**A.** Representative trace of integrated fluorescence intensity of pHrodo Red signal in S305 mutation astrocytes and isogenic controls treated with labelled 0N3R and 0N4R tau isoforms over 48 hours. Error bars = SEM. N = 4, each point is an average of 3 fields of view per well. Linear regression.

**B.** Representative trace of integrated fluorescence intensity of pHrodo Red signal in S305 mutation astrocytes and isogenic controls treated with labelled 1N3R and 1N4R tau isoforms over 48 hours. Error bars = SEM. N = 4, each point is an average of 3 fields of view per well. Linear regression.

**C.** Representative trace of integrated fluorescence intensity of pHrodo Red signal in S305 mutation astrocytes and isogenic controls treated with labelled 2N3R and 2N4R tau isoforms over 48 hours. Error bars = SEM. N = 4, each point is an average of 3 fields of view per well. Linear regression.

*p < 0.05, **p < 0.01, ***p < 0.001, ****p < 0.0001.

**Figure S6. PAM site modification does not alter *MAPT* exon 10 splicing.**

**A.** Sanger sequencing trace indicating V300V PAM site mutation (GUC > GUA). Modified base is highlighted with an asterisk.

**B.** Predicted free energies (kcal) of structures at the Exon 10 – Intron 10 hairpin loop in WT and PAM site mutation V300V sequences. The Exon-Intron junction is indicated with an arrow and the V300V modification is indicated with a red box. The structure of the WT sequence was reported by Lisowiec *et al*.^81^ V300V mutant structure was predicted using RNAstructure.^91^

**C.** Representative RTPCR of *MAPT* Exon 10 minigenes edited with V300V or S305 mutations, expressed in SH-SY5Y cells. Top band is 4R tau, bottom band is 3R tau.

## SUPPLEMENTARY TABLES

*See Excel sheets*

