## Supplementary figures and images for "Development of MAPT S305 mutation models exhibiting elevated 4R tau expression, resulting in altered neuronal and astrocytic function"

### SFig1

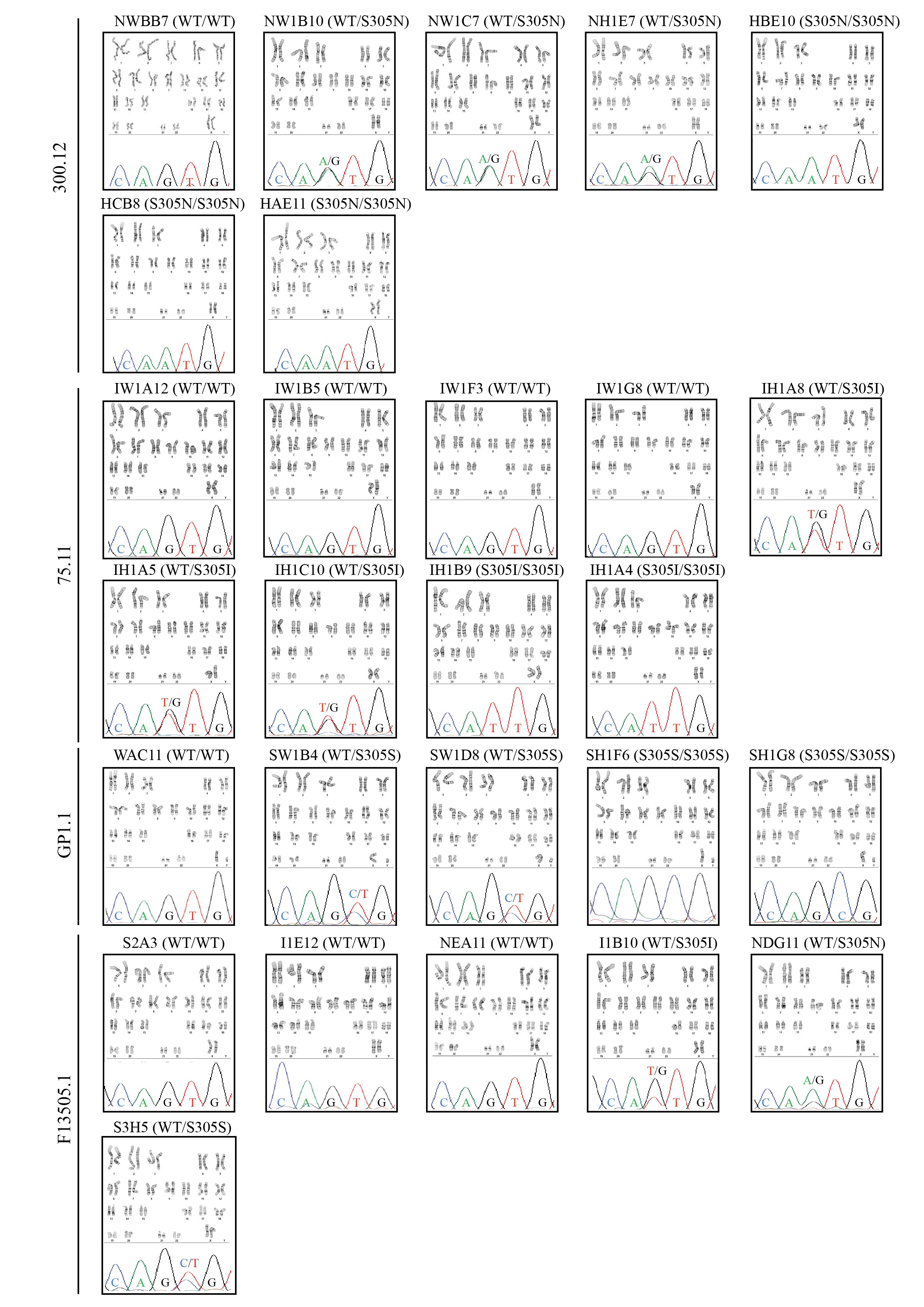

### SFig2

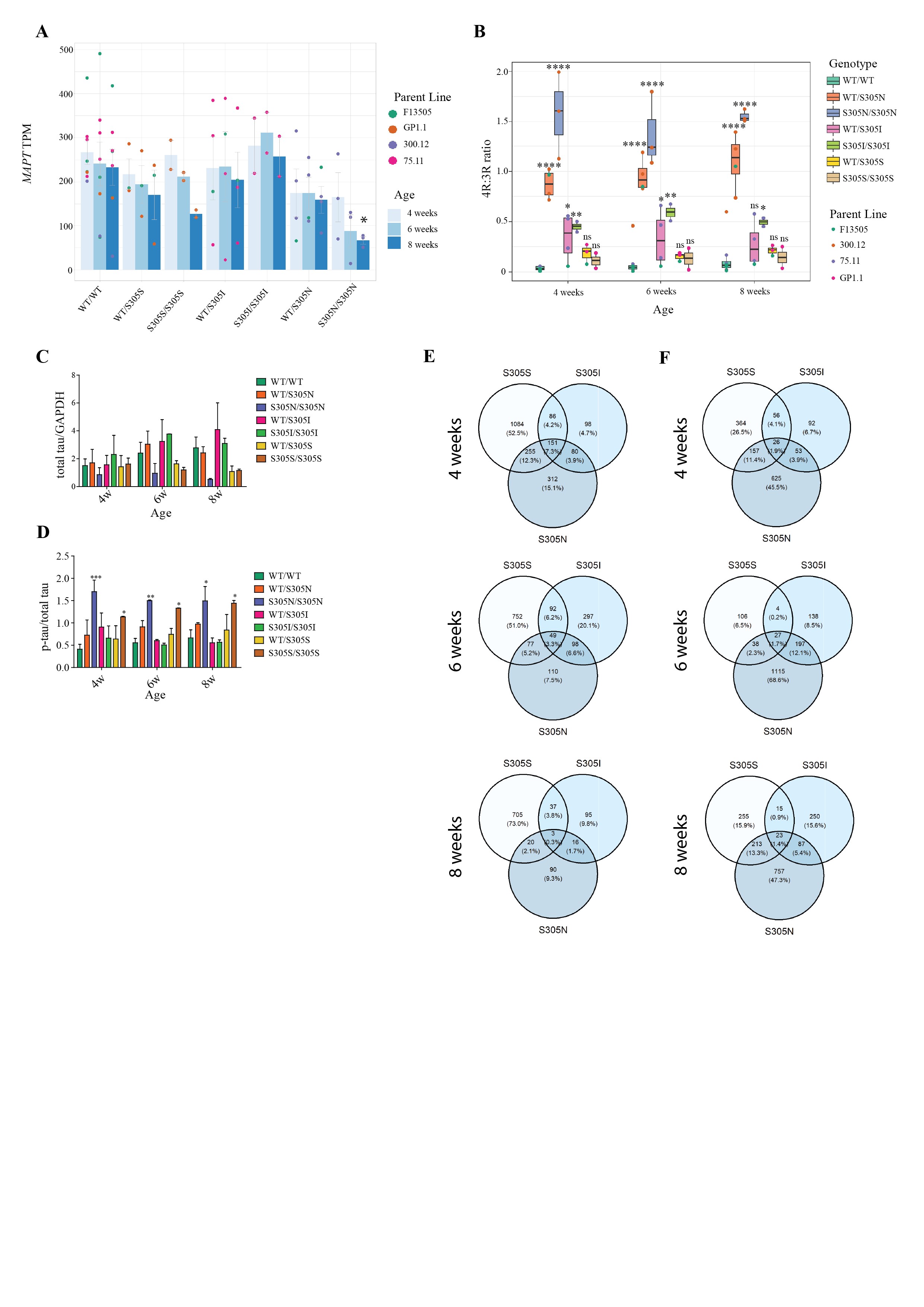

### SFig3

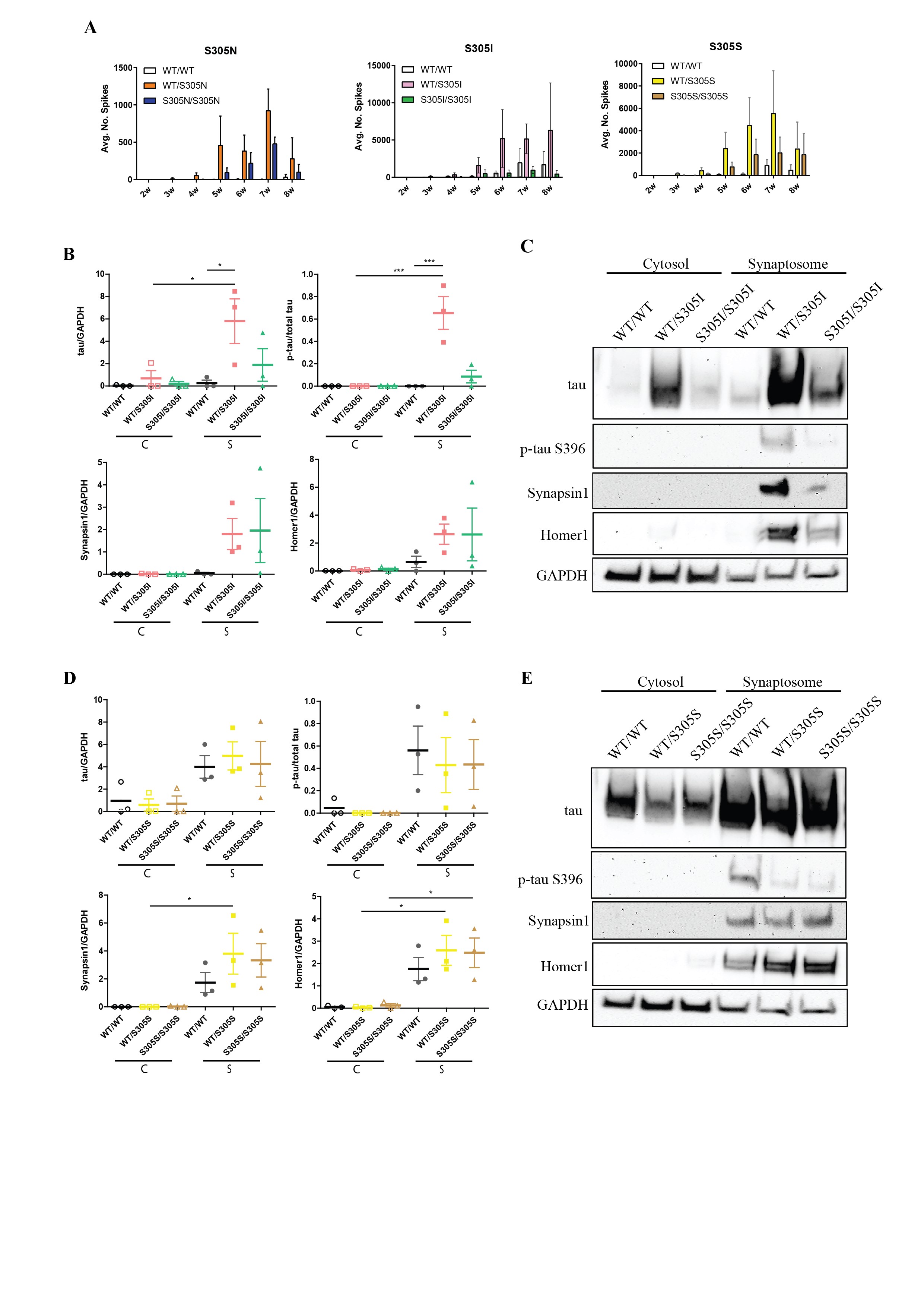

### SFig4

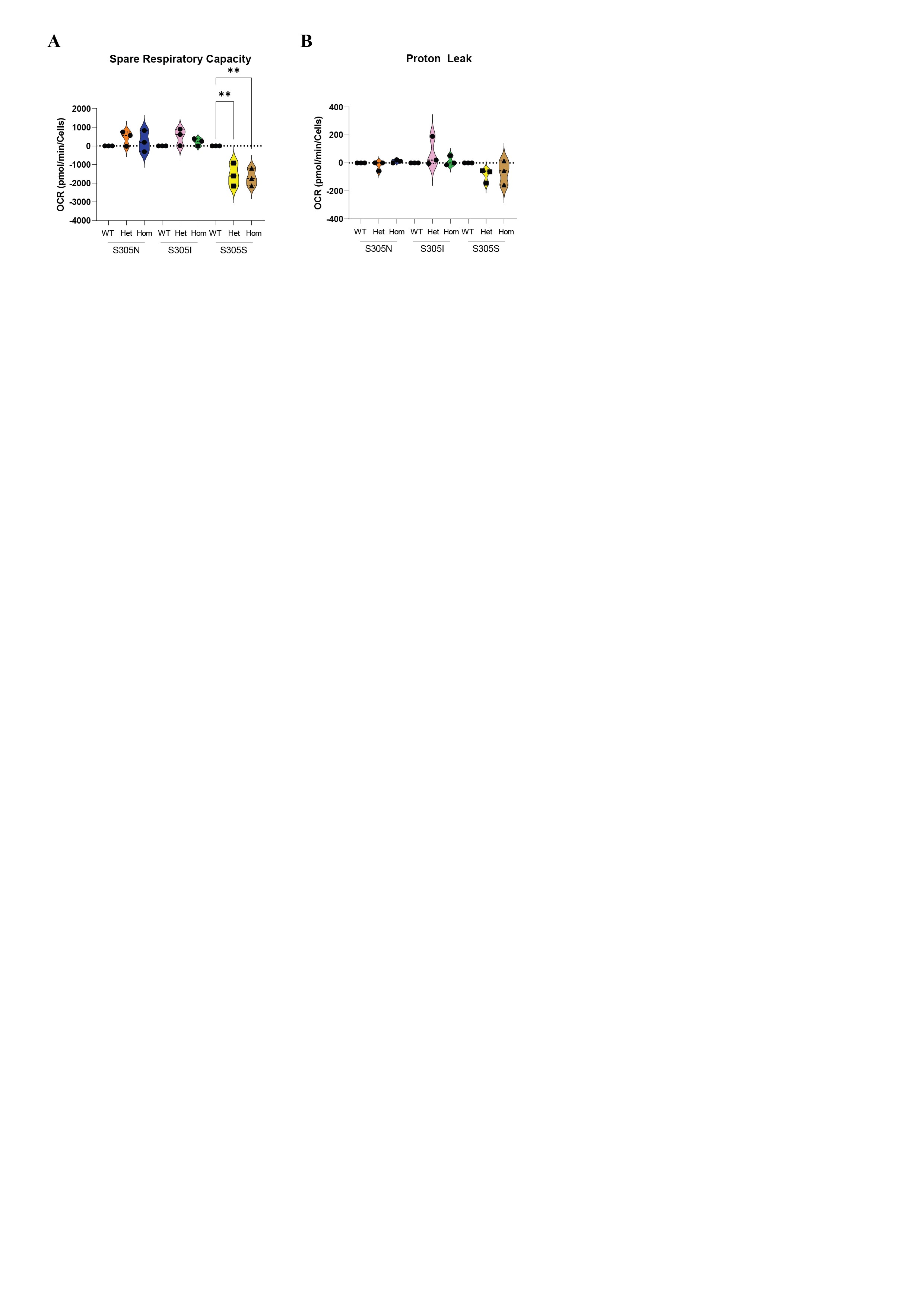

### SFig5

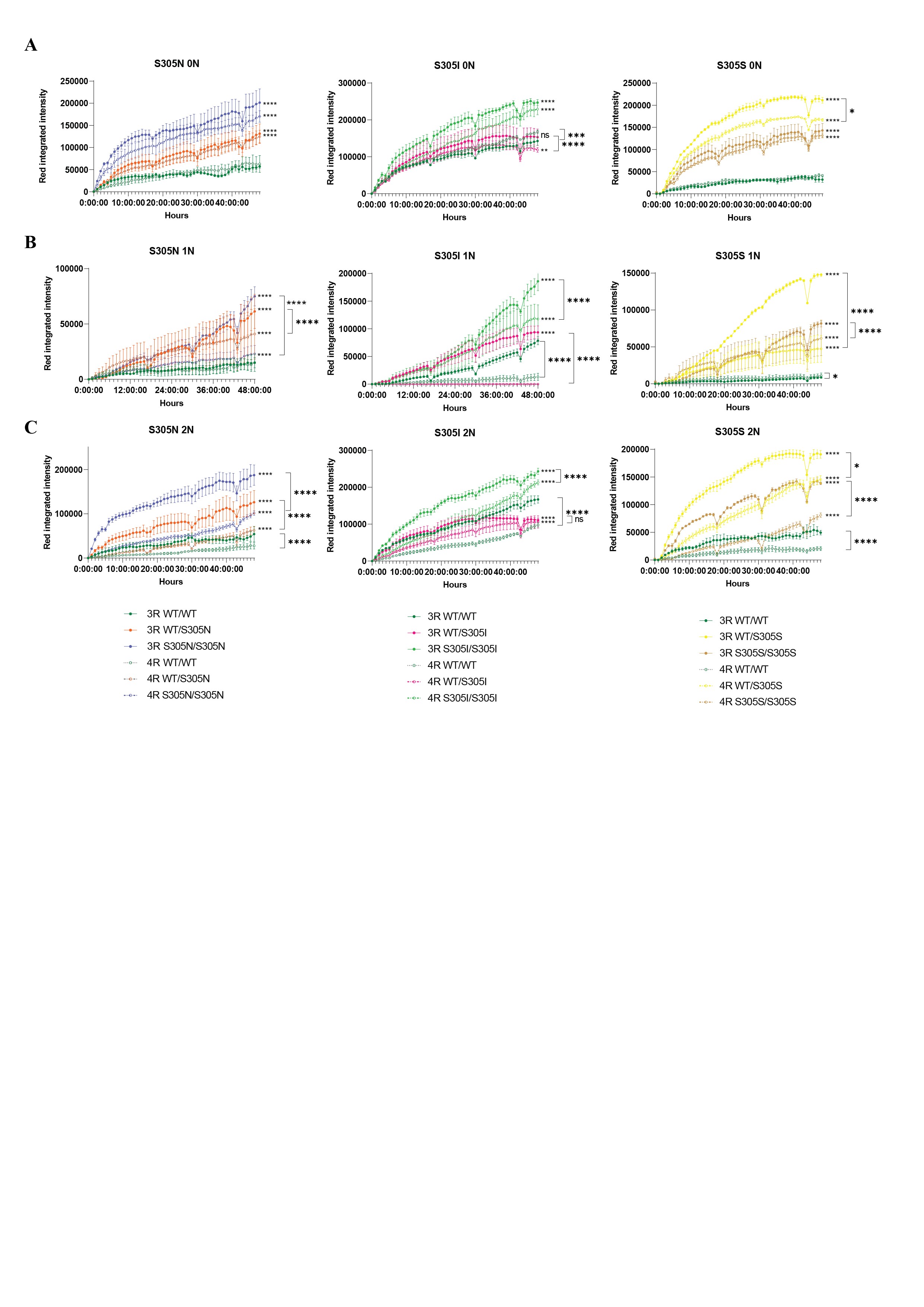

### SFig6

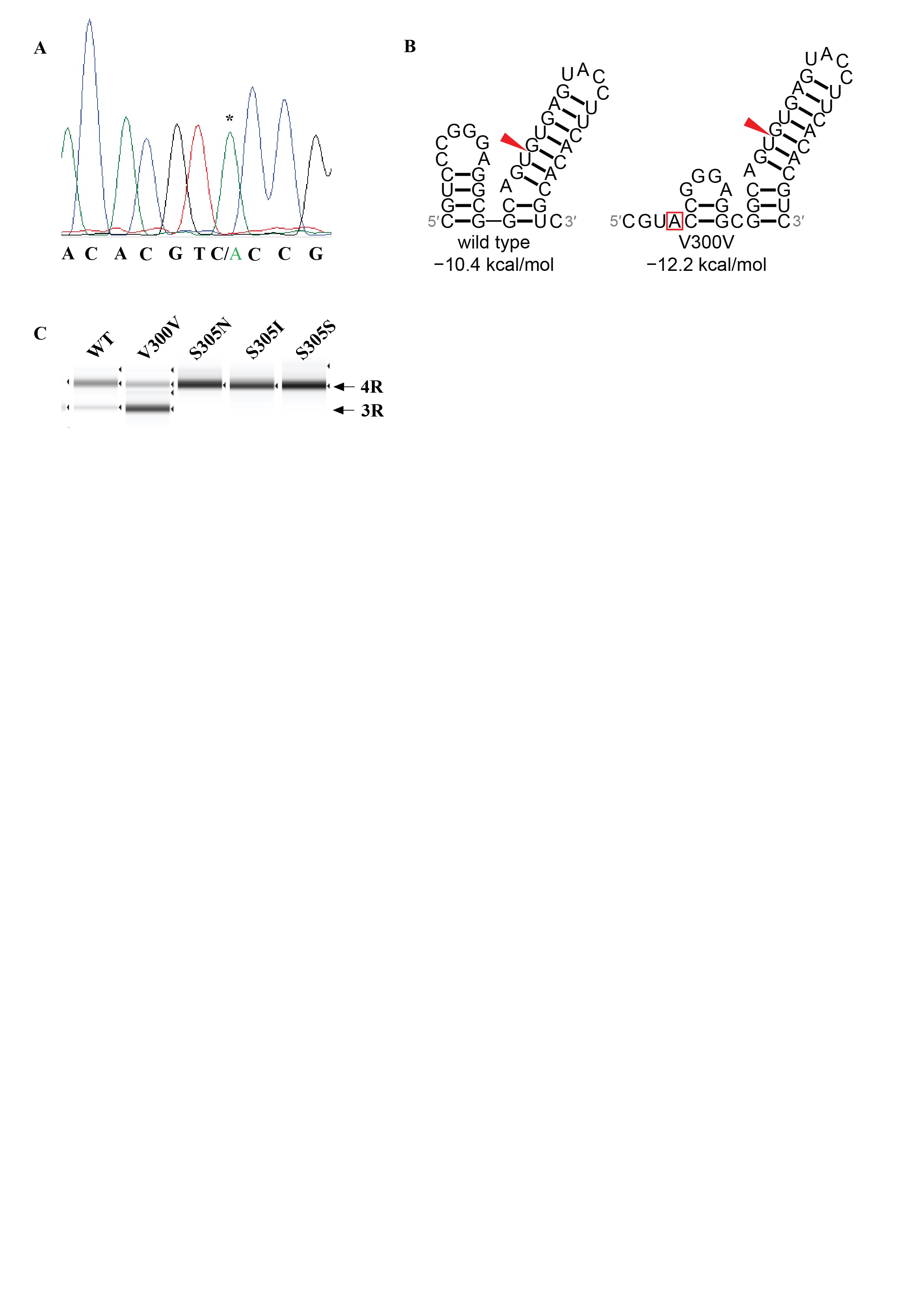
